# Induction of pancreatic neoplasia in the *KRAS*/*TP53* Oncopig

**DOI:** 10.1101/2020.05.29.123547

**Authors:** Pinaki Mondal, Neesha S. Patel, Katie Bailey, Shruthishree Aravind, Michael A. Hollingsworth, Audrey J. Lazenby, Mark A. Carlson

**Affiliations:** Department of Surgery, University of Nebraska Medical Center, Omaha, Nebraska, USA; Department of Surgery and VA Research Service, Nebraska-Western Iowa Health Care System, Omaha, NE, USA; Eppley Institute for Research in Cancer and Allied Diseases, Fred and Pamela Buffett Cancer Center, University of Nebraska Medical Center, Omaha, NE, USA; Department of Pathology, University of Nebraska Medical Center, Omaha, Nebraska, USA; Department of Genetics, Cell Biology and Anatomy, University of Nebraska Medical Center, Omaha, Nebraska, USA

## Abstract

**Introduction:** Five year survival of pancreatic cancer (PC) remains low. Current murine models may not adequately mimic human PC and can be too small for medical device development. A large animal PC model could address these issues. We induced and characterized pancreatic tumors in Oncopigs (transgenic swine with a somatic floxed cassette containing *KRAS*^G12D^ and *TP53*^R167H^).

**Methods:** Oncopigs underwent injection of adenovirus expressing Cre recombinase (AdCre) +/– interleukin 8 (IL-8) into one of the main pancreatic ducts (induction procedure). Subjects were necropsied after ≤10 week, followed by histological analysis, cytokine expression analysis, exome sequencing and transcriptome analysis of resultant tumors.

**Results:** Fourteen Oncopigs underwent the induction procedure; ten (71%) had gross tumor within three weeks, one of these subjects expired suddenly and the other 9 required premature euthanasia secondary to lack of oral intake. At necropsy all of ten of these subjects had gastric outlet obstruction secondary to pancreatic tumor and phlegmon. Two Oncopigs underwent a control injection (no AdCre) and four WT littermates of the Oncopigs underwent AdCre injection without notable effect. Exome and transcriptome analysis of the porcine pancreatic tumors revealed similarity with the molecular signatures and pathways of human PC.

**Conclusion:** Oncopigs with ductal injection of AdCre developed pancreatic tumor in a short period of time with molecular characteristics similar to human PC. While further optimization and validation of this porcine PC model would be beneficial, it is anticipated that this model will be useful for focused research and development of diagnostic and therapeutic technologies for PC.

## INTRODUCTION

Murine models have been used in the preclinical study of pancreatic cancer,^1–4^ including xenografted immunodeficient models, genetically engineered mouse (GEM) models, humanized mice, and *in vivo* edited mice. While murine modeling has produced tremendous advances in the understanding and treatment of cancer in general, the fact remains that only a few percent of drug candidates identified in preclinical studies as potential anti-cancer treatments will be demonstrated to be safe and efficacious in clinical trials, and subsequently obtain FDA approval.^2, 5–7^ Murine models may not accurately predict human biology and response to interventions in all circumstances, secondary to differences in genomic sequence and phenotype. Model inaccuracy may contribute to the above low drug approval rate for potential cancer therapeutics.^4, 8^

Currently there are no validated large animal models for PC, though some proof-of-principle studies in swine have been published.^9–11^ Swine have shown to be effective models in other fields, including trauma, transplantation, cardiovascular disease, and dermatologic conditions.^12, 13^ Swine have greater similarity to humans with respect to size, anatomy, physiology, pathophysiological responses, and coding sequence than do mice.^14, 15^ It therefore is conceivable that swine could have greater accuracy in cancer modeling than mice. Importantly, a porcine PC model would permit research and development of imaging instrumentation, interventional catheters, and other devices suitable for the human PC patient. Such R&D would be difficult if not infeasible in a twenty-gram mouse.

In 2015, a transgenic swine known as the Oncopig Model (OCM) was described,^16^ which utilized a Cre-Lox system to control expression of a somatic cassette containing porcine *KRAS*^G12D^ and *TP53*^R167H^. The expressed KRAS mutant was constitutively activated, while the p53 mutant functioned as a dominant negative. The OCM is the porcine analogue of the KPC mouse.^17^ Previously we attempted induction of PC in the OCM with introduction of relatively low doses of adenovirus expressing Cre recombinase (AdCre) into the OCM pancreatic duct, but we did not obtain tumor.^18^ Herein we report modification of the AdCre injection protocol which successfully generated PC in the OCM. These tumors underwent histologic, exomic, and expressional analysis, with comparison to human PC.

## MATERIALS AND METHODS

### Experimental Subjects and Design

Transgenic Oncopigs (Oncopig Model, or OCM; LSL-*KRAS*^G12D^-IRES-*TP53*^R167H^) and their wild type (WT) littermates were purchased from the National Swine Research and Resource Center (NSRRC) at the University of Missouri Columbia (nsrrc.missouri.edu). The OCM subjects were a hybrid of Minnesota minipigs and domestic pigs. The genotype of each porcine subject was confirmed with PCR upon subject delivery (Fig. S1). Swine were housed two littermates per pen, except for one week of individual housing post-laparotomy (but with contact through the pen grating), in order to prevent wound cannibalism. Swine were fed ad lib with standard hog feed (Purina Nature’s Match® Sow and Pig Complete Feed; www.purinamills.com). The basic experimental design included a ≥1 week acclimatization period after subject delivery to the research facility. Each subject then underwent an induction procedure (laparotomy under general anesthesia; one major survival procedure), followed by observation for up to 3 months.

### Survival Procedure: Tumor Induction

#### Laparotomy and Exposure

A 15 cm upper midline laparotomy incision was made. Abdominal wall retraction was maintained with a self-retaining abdominal (Bookwalter) retractor. The long tongue of the spleen was placed into the right upper quadrant. The small intestine was held inferiorly with laparotomy pads and the self-retaining retractor. The pylorus was identified, grasped, and brought up to the incision. The proximal pancreas could be elevated with the pylorus and proximal duodenum with minimal dissection; this maneuver provided access to both the anterior and posterior surface of the proximal pancreas (see Results and Discussion). The colon typically was lightly adherent to the anterior surface of the pancreas with loose connective tissue, so the colon was mobilized inferiorly from the pancreas with scissors.

#### Injection of Tumor Induction Reagent

There were three basic techniques of injecting the tumor induction reagent into the pancreas: (1) injection into the main pancreatic duct with parenchymal injections; (2) injection into the duct of the connecting lobe of the pancreas; and (3) technique 2 plus parenchymal injections. The induction reagent consisted of AdCre (adenovirus expressing Cre recombinase) at a concentration of 1×10^10^ PFU/100 µL (in saline), with an injection volume of 200 µL. Some subjects (see Results and Discussion) also received porcine IL-8 (5 ng/mL in the same 200 µL volume, mixed in with the AdCre and given as one injection).

#### Technique 1: Main Duct + Parenchyma

After the above exposure of the pancreas, a 3 cm longitudinal enterotomy was made on the anti-mesenteric side of the duodenum, directly opposite to the termination of the duodenal lobe of the pancreas into the duodenal wall, which was the location of the papilla of the main pancreatic duct (MPD). The incised duodenal walls were retracted laterally with silk stay sutures. The MPD then was directly cannulated with a 20-gauge plastic catheter (Angiocath™ IV Catheter; Becton Dickinson) and injected with the induction reagent. Parenchymal injections of the same induction cocktail were also performed in some subjects on the anterior and posterior side of the proximal duodenal lobe of the pancreas (typically two injections on each side, using a 25-gauge needle). Each parenchymal injection site was marked with small dot (1-2 mm wide) of India ink, using a 27-gauge needle. The duodenal enterotomy then was closed in two layers, using a running 3-0 polyglactin 910 suture for the inner row, full thickness, inverting (Connell) technique. This was followed with an outer seromuscular row of interrupted 3-0 silk sutures (Lembert technique).

#### Technique 2: Connecting Lobe

After exposure of the pancreas was obtained, the connecting lobe of the pancreas was identified where it joined the proximal duodenal lobe (see Results & Discussion). The connecting lobe then was doubly clamped and transected about 1 cm inferior to the junction with the duodenal lobe. The proximal stump of the connecting lobe (against the duodenal lobe) was tied off with a 3-0 silk ligature. Using operative loupe (3.5x) magnification, the open end of the primary duct running through the connecting lobe was found in the free cut end of that lobe (see Results and Discussion). This duct was cannulated with a 20-gauge plastic catheter, and the induction reagent was injected into the duct of the connecting lobe. The catheter was withdrawn, and the free end of the connecting lobe then was immediately ligated with 3-0 silk.

#### Laparotomy Closure

The abdominal incision was closed anatomically in layers, with running 3-0 polyglactin 910 in the peritoneum and posterior sheath, running 0-polydioxanone in the anterior aponeurosis, 3-0 polyglactin 910 in the panniculus carnosus, and 4-0 polyglactin 910 in the dermis. Cyanoacrylate glue was applied over the skin incision; no other incisional dressing was applied. The animal’s recovery from anesthesia was monitored until the subject was awake and mobile. Subjects were given half feeds on post-induction day 1 and were placed back on ad lib feeds by day 2.

#### Additional Methods and Materials

Conduct of animal experiments was guided by the Animal Research: Reporting of In Vivo Experiments (ARRIVE) standards (Fig. S6). See Supplemental Material for description of animal welfare concerns, animal numbers and randomization, anesthesia and analgesia, euthanasia, histology, serum testing, sequencing and statistical analysis, and other Methods (Fig. S2).

## RESULTS

### Relevant Anatomy of the Porcine Pancreas

In humans, the pancreas has been labeled with the somewhat arbitrary anatomical regions of “head” (adjacent to the duodenum and to the right of the portomesenteric vein), “neck” (overlying the portomesenteric vein), “body” (to the left of the portomesenteric vein), “tail” (near the splenic hilum), and a variably-present uncinate process which comes off the inferior side of the pancreatic head.^19^ In the pig, however, there are three fairly distinct lobes^20^ which interconnect to form the pancreas: the duodenal, splenic, and connecting lobes (Fig. 1A). As described later, this tri-lobar configuration becomes critically important for the porcine pancreatic cancer model. The duodenal lobe of the porcine pancreas (Fig. 1A-B) is an elongate structure situated inferiorly to the pylorus and gastric antrum. The proximal end of the duodenal lobe tapers to a termination on the mesenteric border of duodenal C-loop, 10-15 cm distal to the pylorus. At this point the main pancreatic duct traverses the duodenal wall and opens into the duodenal lumen via the pancreatic papilla. The splenic lobe of the porcine pancreas (Fig. 1A-B) is another elongate structure which initiates at the distal end of the duodenal lobe and terminates near the splenic hilum.

**Fig. 1.**
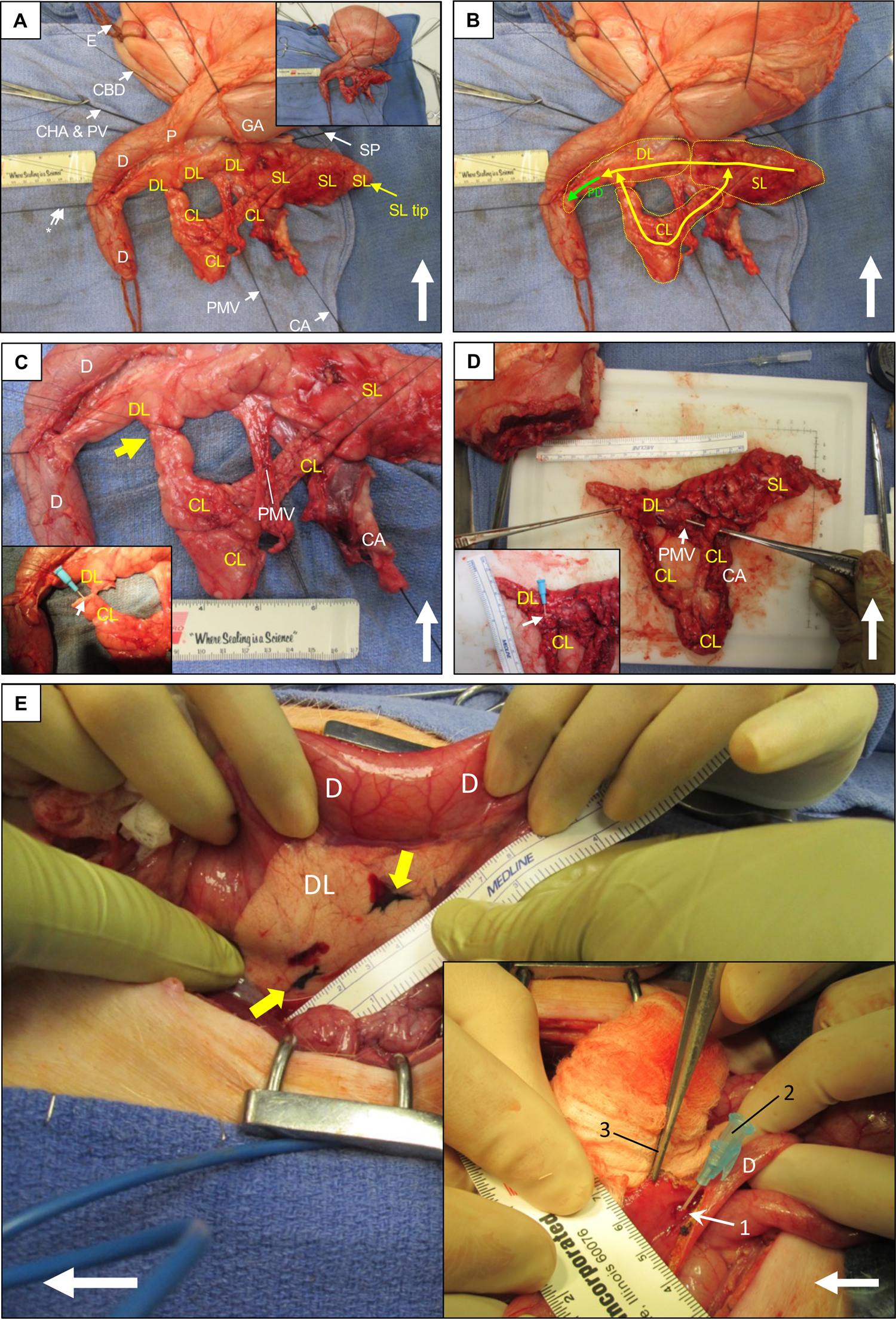
Porcine pancreatic anatomy and injection techniques. (**A**) Anterior aspect; top = cephalad. DL = duodenal lobe; CL = connecting lobe; SL = splenic lobe. *Double arrow suture = main pancreatic duct entering into duodenum via the pancreatic papilla (not visible), CA = celiac artery; CBD = common bile duct; CHA & PV = common hepatic artery and portal vein (not visible); E = esophagus; GA = gastric antrum; P = pylorus; PMV = portomesenteric vein emerging from underneath pancreas; SP = splenic pedicle. Inset: zoom out view. (**B**) Same specimen as panel **A**. Dashed yellow boundaries = pancreatic lobes. Solid thick yellow lines = approximate course of pancreatic ductal system. Intersections of the DL, SL, and CL form a continuous ductal loop; if interrupted, system decompresses retrograde into the main pancreatic duct (PD, green arrow). (**C**) Same specimen, zoom in. Yellow arrow = CL transection for ductal injection. Inset: CL has been transected; 22g plastic catheter inserted (small white arrow) into CL duct (Technique 2 of tumor induction). (**D**) Similar dissection in another subject, demonstrating circular DL-SL-CL lobar configuration. Inset: cannulization of transected CL duct with a 22g plastic catheter. (**E**) Pancreatic parenchymal and main duct injection (Technique 2). Duodenal lobe (DL) inked (yellow arrows) for parenchymal AdCre injection. Inset: PD accessed through a duodenotomy just opposite to the insertion of the PD; pancreatic papilla (1) then cannulated with 22g catheter (2) for AdCre injection; Debakey forceps (3) retracts duodenal wall. White arrows = cephalad.

The connecting lobe of the porcine pancreas (Fig. 1A-D) is a structure unique to the pig and has no equivalent in humans. The connecting lobe is a U-shaped structure which connects the mid-portion of the duodenal lobe to the mid-portion of the splenic lobe. This tri-lobar configuration forms a continuous ring of the porcine pancreas, within which runs a continuous circular ductal system (Fig. 1B). This circular configuration is reminiscent of the arterial Circle of Willis which is located at the base of the human brain. The main pancreatic duct of the pig exits from this circular ductal system within the duodenal lobe (Fig. 1B), and continues proximally to enter the duodenum, as described above. This circular anatomy of the porcine pancreas affords a unique surgical opportunity, in that the ring can be safely interrupted for an intervention (e.g., ductal injection, Fig. 1C-D). After such an interruption, the ductal system will be capable of retrograde decompression away from the interruption point, and not be obstructed.

### Connecting Lobe Injection in WT Pigs

As mentioned in the Introduction, we had previously attempted tumor induction^18^ in the OCM (N = 5) but did not obtain gross tumor after 4 months of observation (Fig. S3). For those induction procedures, AdCre (the induction reagent) was injected into the main pancreatic duct and parenchyma of the duodenal lobe (Technique 1 in the Methods) at a dose of 1 x 10^8^ pfu. Some histologic abnormalities were noted (proliferative ductal lesions), but none of these appeared to be cancerous. Of note, Schook et al.^11^ observed a diffuse nodular thickening of the main pancreatic duct which histologically appeared to be adenocarcinoma in one OCM subject, 12 months after intraductal injection of 4 x 10^9^ pfu of AdCre. We hypothesized that if we increased our dose of AdCre that was injected into the pancreatic duct, then we should obtain ductal cancer in the OCM.

One issue that we noted with our previous group of five OCM subjects^18^ (Fig. S3) was that after injection of AdCre into main pancreatic duct, the continual flow of pancreatic secretion rapidly flushed the injectate out of the papilla and into the duodenal lumen. This flushing rapidly diluted the AdCre, likely decreasing its efficacy. In order to prevent this flushing, the pancreatic duct would need to be ligated. However, ligation of the main pancreatic duct would obstruct the entire ductal system (Fig. 1A-B), which in swine produces pancreatic exocrine insufficiency.^21^ We hypothesized that we could instead transect the isthmus region of the proximal connecting lobe (Fig. 1A-D), cannulate and inject the pancreatic duct to the connecting lobe, and then immediately ligate the duct ends with impunity secondary to the circular ductal anatomy of the porcine pancreas (Fig. 1B). Even with the connecting lobe interrupted as above, the entire pancreas could still decompress into the main pancreatic duct (Fig. 1B) after such a maneuver. This ductal injection and ligation would minimize loss of the injectate, which itself would be in contact with the ductal epithelium for a longer duration than had the duct been left open.

In order to determine whether interruption of the isthmus portion of the connecting lobe (along with cannulation of the duct within the connecting lobe) would feasible and well-tolerated, we undertook this procedure in WT domestic pigs (N = 5; 3–4 months, 30-45 kg; see Table S1 for descriptive data of porcine subjects), injecting saline distally into the connecting lobe, with ligation of both cut ends of that lobe (i.e., induction technique 2 in the Methods). This technique effectively interrupted the circular configuration of the porcine pancreas. At operation, the duct within the transected connecting lobe was small (≤1 mm) but could accommodate insertion of a 20-gauge plastic catheter (Fig. 1B) and flushed easily with 1.0 mL of saline. All five pigs tolerated the connecting lobe injection procedure without difficulty. All pigs underwent necropsy one month after operation, and there was no evidence of pancreatic inflammation, exocrine insufficiency, or other adverse effect from ductal interruption (data not shown).

### Induction of Tumor in OCM Subjects: Technique 1

We elected to give a higher dose of AdCre (2 x 10^10^ pfu) with the current experiments than what we used before (1 x 10^8^ pfu),^18^ i.e., >100-fold increase in the dose of AdCre. We also elected to administer IL-8 as an adjunct with the AdCre; this cytokine has been shown to mobilize the adenoviral receptor to the luminal membrane of epithelial cells, thereby enhancing viral entry.^22, 23^ The intention of these two changes (high AdCre dose and IL-8) was to increase the chance for successful pancreatic tumor induction in the OCM.

The first subject (ID. 468, Table 1) underwent induction with technique 1 (i.e., AdCre + IL-8 injection into the main pancreatic duct + injections into the parenchyma of the duodenal lobe); see Fig. 2A. This subject died unexpectedly over a weekend 17 days after the induction procedure. The carcass was refrigerated, and the necropsy was performed 72 hours later. The subject had gastric perforation with extensive peritoneal soilage (Fig. S4A). The cause of the gastric perforation was gastric outlet obstruction, which in turn was caused by a peripancreatic phlegmon (Fig. S4B). Unfortunately, reasonably intact tissue could not be obtained for histology secondary to the prolonged interval between death and necropsy. Our initial suspicion was that subject 468 developed severe pancreatitis, but not necessarily tumor. The second subject (ID. 469, Table 1) underwent a similar tumor induction procedure and underwent euthanasia/necropsy on day 73 (planned endpoint) after an uncomplicated course. There was no gross or microscopic evidence of tumor.

**Fig. 2.**
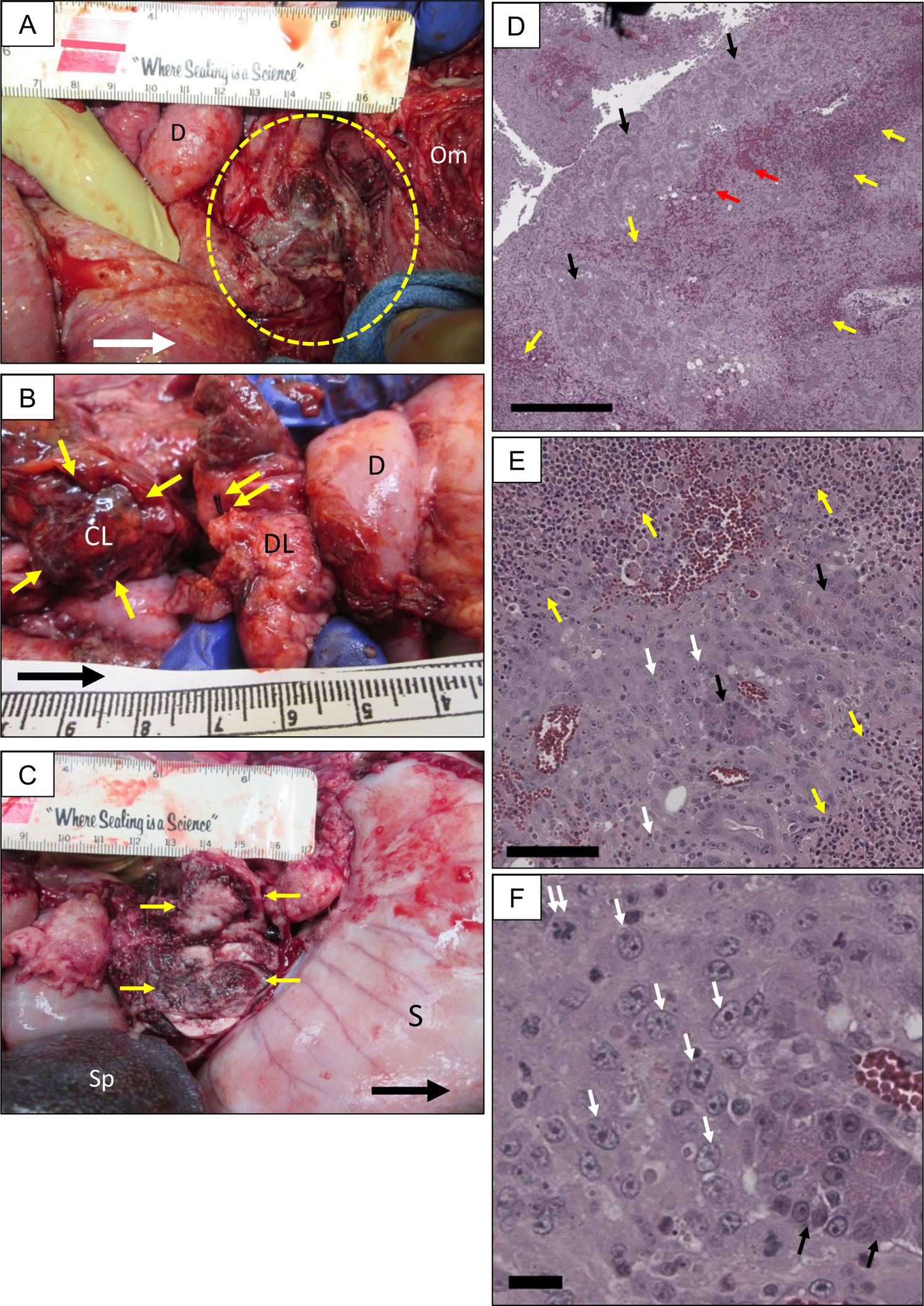
Induction of OCM pancreatic tumor. (**A**) OCM necropsy <3 wk after the tumor induction procedure; dashed yellow line = pancreatic phlegmon (contained tumor) at location of AdCre injection. D = duodenum; Om = omentum. Scale = cm; large arrows = cephalad. (**B**) Another OCM necropsy <3 wk after tumor induction procedure. Silk suture (double yellow arrow) = ligated proximal end of connecting lobe (CL) at intersection with duodenal lobe (DL). Single yellow arrows = distal CL remnant (site of tumor). (**C**) Third OCM necropsy <3 wk after tumor induction. Transection of CL phlegmon (yellow arrows) demonstrated firm nodular mass (tumor on pathology). Subject had typical gastric outlet obstruction (distended stomach, S); Sp = spleen. (**D**) Low power view (H&E) of CL injection site (pancreatic phlegmon). Cords of inflammatory cells (yellow arrows) intermingled with hemorrhage (red arrows), with residual acini (black arrows). Bar = 500 µm. (**E)** Higher power view of phlegmon from panel **D**. Sheets of tumor cells were apparent (white arrows), intermingled with residual acinar structures. Bar = 100 µm. (F) High power view from panel **E**; individual tumor cells indicated with white arrows; double arrow = mitotic figure. Bar = 20 µm.

**Table 1.**
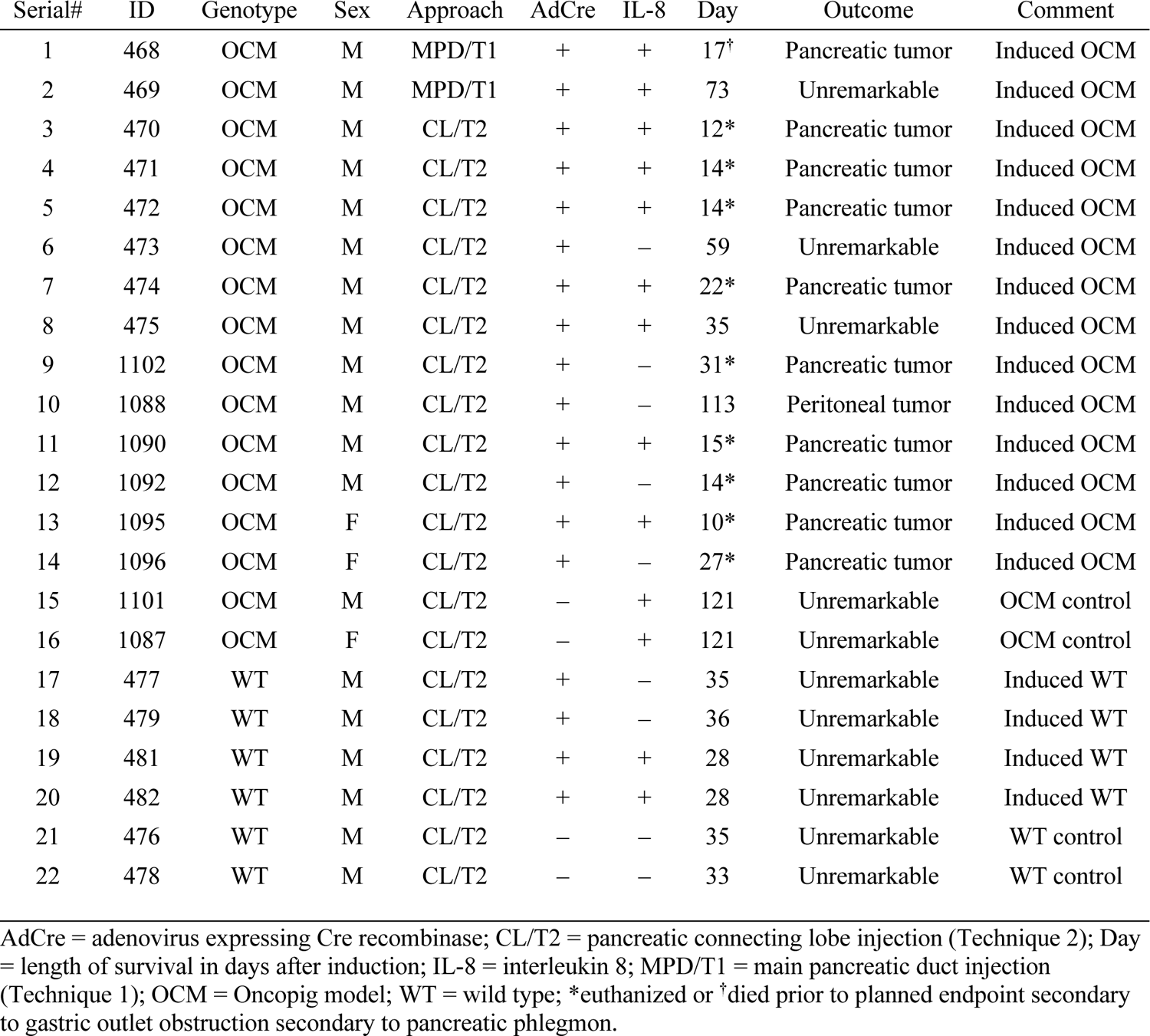
Tumor induction method and outcome in OCM and WT pigs.

| Serial# | ID | Genotype | Sex | Approach | AdCre | IL-8 | Day | Outcome | Comment |
| --- | --- | --- | --- | --- | --- | --- | --- | --- | --- |
| 1 | 468 | OCM | M | MPD/T1 | + | + | 17<sup>†</sup> | Pancreatic tumor | Induced OCM |
| 2 | 469 | OCM | M | MPD/T1 | + | + | 73 | Unremarkable | Induced OCM |
| 3 | 470 | OCM | M | CL/T2 | + | + | 12* | Pancreatic tumor | Induced OCM |
| 4 | 471 | OCM | M | CL/T2 | + | + | 14* | Pancreatic tumor | Induced OCM |
| 5 | 472 | OCM | M | CL/T2 | + | + | 14* | Pancreatic tumor | Induced OCM |
| 6 | 473 | OCM | M | CL/T2 | + | – | 59 | Unremarkable | Induced OCM |
| 7 | 474 | OCM | M | CL/T2 | + | + | 22* | Pancreatic tumor | Induced OCM |
| 8 | 475 | OCM | M | CL/T2 | + | + | 35 | Unremarkable | Induced OCM |
| 9 | 1102 | OCM | M | CL/T2 | + | – | 31* | Pancreatic tumor | Induced OCM |
| 10 | 1088 | OCM | M | CL/T2 | + | – | 113 | Peritoneal tumor | Induced OCM |
| 11 | 1090 | OCM | M | CL/T2 | + | + | 15* | Pancreatic tumor | Induced OCM |
| 12 | 1092 | OCM | M | CL/T2 | + | – | 14* | Pancreatic tumor | Induced OCM |
| 13 | 1095 | OCM | F | CL/T2 | + | + | 10* | Pancreatic tumor | Induced OCM |
| 14 | 1096 | OCM | F | CL/T2 | + | – | 27* | Pancreatic tumor | Induced OCM |
| 15 | 1101 | OCM | M | CL/T2 | – | + | 121 | Unremarkable | OCM control |
| 16 | 1087 | OCM | F | CL/T2 | – | + | 121 | Unremarkable | OCM control |
| 17 | 477 | WT | M | CL/T2 | + | – | 35 | Unremarkable | Induced WT |
| 18 | 479 | WT | M | CL/T2 | + | – | 36 | Unremarkable | Induced WT |
| 19 | 481 | WT | M | CL/T2 | + | + | 28 | Unremarkable | Induced WT |
| 20 | 482 | WT | M | CL/T2 | + | + | 28 | Unremarkable | Induced WT |
| 21 | 476 | WT | M | CL/T2 | – | – | 35 | Unremarkable | WT control |
| 22 | 478 | WT | M | CL/T2 | – | – | 33 | Unremarkable | WT control |

AdCre = adenovirus expressing Cre recombinase; CL/T2 = pancreatic connecting lobe injection (Technique 2); Day = length of survival in days after induction; IL-8 = interleukin 8; MPD/T1 = main pancreatic duct injection (Technique 1); OCM = Oncopig model; WT = wild type; \*euthanized or <sup>†</sup>died prior to planned endpoint secondary to gastric outlet obstruction secondary to pancreatic phlegmon.

After these two subjects we hypothesized that injecting reagents into the main pancreatic duct was not allowing efficient AdCre action secondary to the flushing phenomenon described above. We then attempted connecting lobe injection (technique 2 in the Methods) for subsequent tumor inductions. We also decided to forgo the concomitant parenchymal injections, secondary to the concern that parenchymal injection would increase exposure of a wide variety of cell types (pancreatic ductal cells, islet cells, fibroblasts, endothelial cells) to AdCre and possible transformation. Pleomorphic or sarcomatous tumor induction with non-directed AdCre injection in the OCM has been observed by Schook’s group.^11, 16^ We reasoned that injection of AdCre directly into a pancreatic would should keep viral exposure mostly confined to the epithelial cells lining the lumen of the duct, and therefore produce tumor that would be (mostly) an adenocarcinoma.

### Induction of Tumor in OCM Subjects: Technique 2

Pancreatic tumor induction with AdCre injection into duct of the connecting lobe (technique 2) was attempted in 12 OCM subjects (Table 1). Only two of these subjects (ID 473 and 475, Table 1) lived to their planned euthanasia data without incident, and had no remarkable findings at gross necropsy nor microscopic analysis. Nine of the remaining subjects (75%) had onset of lethargy and decreased feeding at 10-15 days after the induction procedure, with progressive decline that necessitated unplanned euthanasia at a mean of 18 ± 7 days (range 10-31). The gross findings at necropsy in these nine subjects was similar to that in OCM no. 468 above (technique #1; unexpected death): a peripancreatic phlegmon producing gastric outlet obstruction (Fig. 2B, Fig. S4C). In each case the phlegmonous process had obliterated the region in and around the proximal pancreas; in some cases, there was gross liquefaction. It was difficult to discern anatomical landmarks around the phlegmon. In addition, six of the subjects with tumor had a varying number of extrapancreatic intraabdominal nodules (<1 cm) studding the surface of various structures (Fig. S4D), including the omentum, mesentery, diaphragm, liver, spleen, parietal peritoneum, stomach, and intestines.

The initial concern was that AdCre injection into the connecting lobe of the OCM subjects was inciting severe pancreatitis that produced gastric outlet obstruction which ultimately was fatal. However, upon cutting into the phlegmon of these subjects there was a firm, somewhat pale core which appeared to be tumor (Fig. 2C). H&E microscopy of slices taken from the core of the peripancreatic phlegmon from all nine subjects that underwent premature euthanasia demonstrated abundant tumor cells (Fig. 2D-F) with large nuclei and prominent nucleoli. Tumor cell morphology varied from spindle-shaped to rounded. Sheets of tumor cells were interspersed with normal appearing pancreatic acini and the occasional islet, along with areas of hemorrhage and liquefaction necrosis. Areas of tumor were often surrounded by areas of activated lymphocytes (Fig. 2D-E). In addition, there were numerous tumor-infiltrating lymphocytes and macrophages, and occasionally neutrophils and eosinophils. In some subjects there were numerous multi-nucleated giant cells which appeared to be engulfing vacuoles of necrotic material.

Immunohistochemistry of tumor samples from the peripancreatic phlegmon stained positive for mutant KRAS^G12D^ and mutant p53 (Fig. 3A). Quantification of mutant KRAS^G12D^ and mutant p53 staining demonstrated cellular positivity rates in tumor of ∼70% and ∼40%, respectively, while nontreated OCM pancreas had positivity rates close to zero (Fig. 3B). Tumor sections also had increased Ki-67 and Alcian blue staining compared to control sections (Fig 3C, D). These data suggested that OCM pancreatic tumors were more proliferative and had increased acidic mucins and hyaluronic acid deposition into the extracellular matrix, which is consistent with human pancreatic cancer.^24–26^ Tumor sections also stained positive for CK19, vimentin and CD31 markers (Fig. 3E), which suggested a mixed epithelial phenotype, possibly with epithelial to mesenchymal transition. Quantification of CK19 staining showed no significant difference between tumor and control samples (data not shown).

**Fig. 3.**
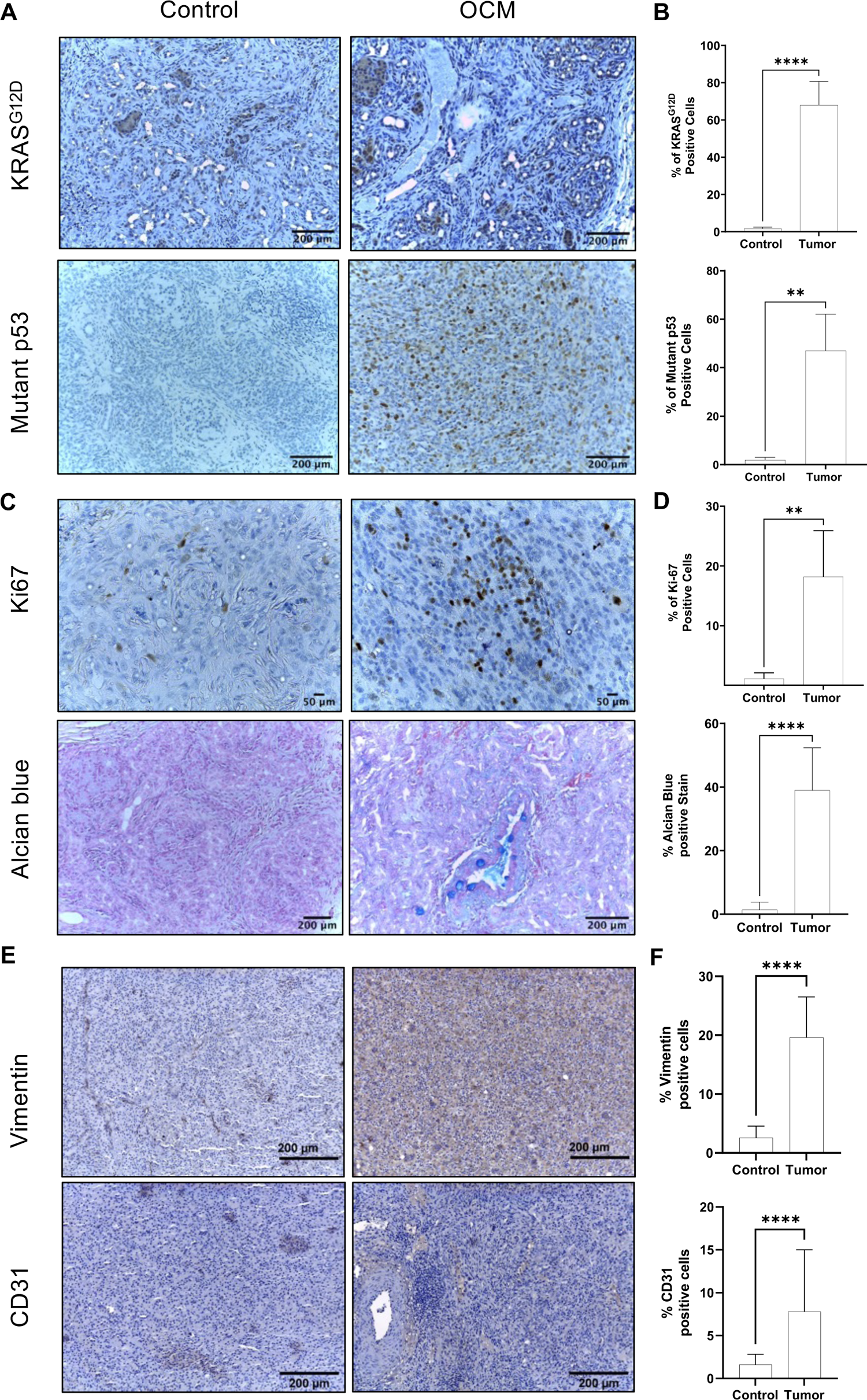
Immunohistochemistry of tumor-associated proteins and polysaccharide staining in pancreatic tumor from AdCre-induced Oncopigs *vs*. pancreas from WT littermates. Expression and quantification of KRAS and p53 mutants. (**A**-**B**) and Ki-67 and Alcian blue (**C**-**D**) and Vimentin and CD31 **(E-F)**. Each bar represents the mean ± SD of four microscopic fields captured from each subject (three subjects each, induced OCM vs. WT); **p <0.01, ****p <0.0001.

The gross appearance of extrapancreatic nodules that were noted in some OCM subjects (Fig. S4) was consistent with “drop” or contact metastases, being only present on the surface of the involved organ or tissue, and mostly inferior and ventral to the phlegmon. There was no evidence of nodule formation within the body of an organ, such as the liver (as is frequently the case in clinical metastatic PC^27^). Histologic evaluation of these extrapancreatic nodules revealed them to be dedifferentiated tumor with a prominent component of activated lymphocytes, both around and within the tumor (data not shown). In some cases, these tumor implants showed evidence of invasion into the underlying normal tissue, with disruption of the peritoneum and underlying connective tissue capsule covering the normal tissue (data not shown).

The remaining, as yet undescribed subject (ID. 1088, Table 1) in this group of 12 OCM with ductal AdCre injection underwent unplanned euthanasia on day 113 secondary to respiratory distress. At necropsy there was gross evidence of acute pneumonia; on histology, the pulmonary alveoli were filled with fluid and acute inflammatory cells. Blood work at necropsy showed an elevated WBC. There also were several nodules (≤1 cm) on the surface of the anterior stomach, but the pancreas was unremarkable. Histologic evaluation of the gastric nodules demonstrated dedifferentiated tumor, similar to the above extrapancreatic nodules. There also were sheets of activated lymphocytes present on H&E sections of the liver, spleen, and lung. The cause of death was deemed to be pneumonia.

### Control Injections in OCM and WT Pigs

Injections of tumor-induction agents (AdCre ± IL-8) into the connecting lobe (i.e., technique 2) of WT littermates (N = 6; Table 1) of the OCM subjects were performed in order to determine whether the procedure or the induction agents would induce any pathology in the background strain of the OCM. All WT subjects tolerated the injection without difficulty, and there were no remarkable findings at necropsy. H&E evaluation of the connecting lobe of the pancreas (i.e., AdCre injection site) in these subjects revealed minimal chronic inflammation, but otherwise was unremarkable (data not shown). Immunohistochemical staining of Ki-67 and Alcian blue was lower in the WT pancreas compared to OCM tumor (Fig. 3C, D).

In order to determine whether some condition unique to the transgenic Oncopig (i.e., not the tumor) was producing severe pancreatitis after the connecting lobe procedure, two OCM subjects underwent IL-8 injection only into the connecting lobe (ID. 1101 and 1087, Table 1). These two subjects tolerated the procedure without difficulty and had no remarkable findings at necropsy. Immunohistochemistry of these two OCM controls showed similar staining to WT controls with AdCre injection into the connecting lobe, including positive staining for CK19, vimentin and CD31 (data not shown). The results of the control injections from both WT and OCM subjects suggested that the severe inflammatory response noted in AdCre-treated OCM subjects was secondary to tumor formation, and not some other response to the pancreatic injection procedure.

### Effect of IL-8 Co-Administration

Sorting the data in Table 1 based AdCre injection with or without IL-8 co-administration revealed pancreatic tumor in 7/9 OCM subjects with IL-8 versus 3/5 subjects without IL-8 (p > 0.48, chi-square), with euthanasia at 15 ± 4 and 24 ± 9 days, respectively (p > 0.2, unpaired t-test).

### Serum Cytokine Levels

Some inflammatory cytokines have been shown to be elevated in patients with pancreatitis or pancreatic cancer, and have been implicated in the development and progression of the latter.^28^ Serum cytokine array analysis performed on three tumor-bearing OCM subjects demonstrated increased expression of IL-1β, IL-6, IL-8 and IL-10 at necropsy compared to pre-induction serum levels (Fig. S5A); the pre-induction vs. necropsy levels of other cytokines in the array were not significantly different for subjects with tumor. The changes in the pre-induction vs. necropsy levels for the above four cytokines in three WT control subjects were less consistent (Fig. S5B), with upward, downward, or no trends being evident. However, statistical comparison of the relative change (pre-induction to necropsy) of each cytokine was not different in the OCM *vs*. WT subjects (Table S2). Pre-induction vs. necropsy levels for the other cytokines in the array not shown in Fig. S5B were not significantly different for WT subjects. The baseline (pre-induction) level of each cytokine shown in Fig. S5B was not different in the OCM *vs*. WT subjects (Table S2).

### Gene variants in OCM pancreatic tumors associated with human PDAC

Exome analysis of four Oncopig pancreatic tumors demonstrated insertions and deletions in all chromosomes in all four tumors (Fig. 4A). Functional variations with high prediction to be able to change protein expression were identified through The Ensemble Variant Effect Predictor.^29^ These high-impact variations on protein expression were present in all the tumors and throughout the exome (Fig. 4B). *KRAS,* the important gene altered in human pancreatic cancer, showed two different types of alterations: deletions and alteration at a known SNP position (Table S3). The exome analysis successfully detected the presence of *KRAS ^G12D^* mutation in the Oncopigs (Table S3). Another pancreatic cancer associated gene in humans, *TP53* showed two intronic variants in three OCM tumors that are in a known SNP position (Table S3). One mutation in only one pig tumor was found in *SMAD4* (Table S3).

**Fig. 4.**
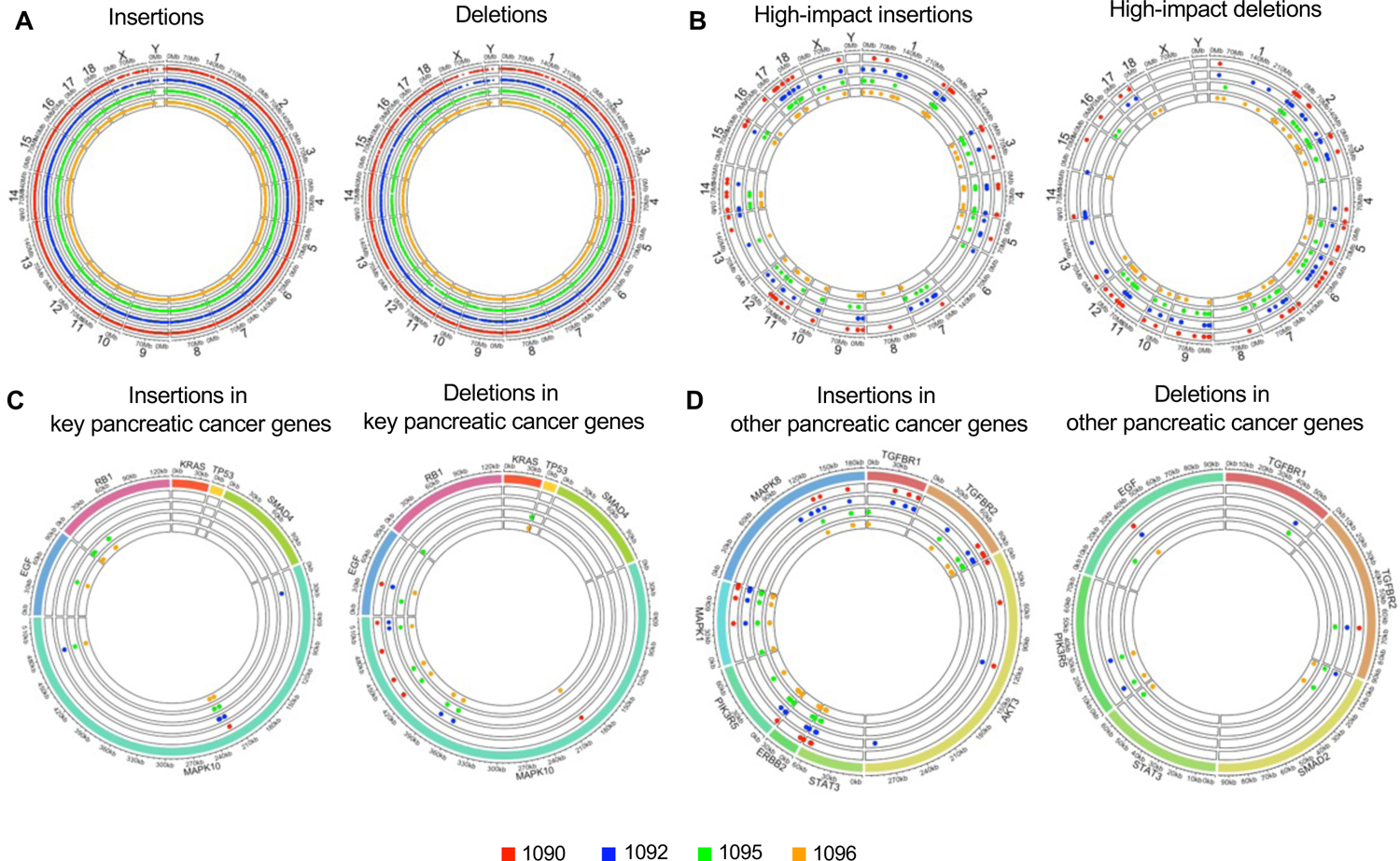
Exome insertions and deletions in OCM tumors. All displayed maps compare exome sequence data between a group of four OCM pancreatic tumors (subjects 1090, 1092, 1095, and 1096 from Table 1) *vs*. normal porcine pancreas (i.e., Oncopig pancreas without AdCre injection); indels (insertions and deletions) refer to tumor exome with respect to normal. (**A**) Exome-wide map of all indels. (**B**) Indels predicted to have an effect on protein expression. (**C**) Indels involving key genes in human PC. (**D**) Indels involving other genes related to human PC.

Apart from these three common genes of human pancreatic cancer, we identified other commonly associated genes with human PDAC which were found to be altered by insertions, deletions, and functionally relevant variations (Table S4 and Fig. 4D). Deletions found in three out four pig tumors in *CCND1*, *CASP9* and in all samples for *ARHGEF6* (Table S4). Insertions and deletions were mostly found in *MAPK10* gene, with occasional presence in *EGF* and *RB1* (Fig. 4C-D). Together, these preliminary findings suggest that porcine pancreatic tumors showed similar mutational landscape with human PDAC.

### Similarities of gene expression in OCM pancreatic tumor vs. human PDAC

The transcriptome of OCM pancreatic tumors was compared with pancreas of wild type littermates. Known human PDAC genes, including *KRAS* and *TP53*, were over-expressed in OCM tumors (Fig. 5A and Tables S3, S4, and S5). Genes associated with epithelial to mesenchymal transition (EMT) pathways,^30^ including *MMP3* (matrix metalloproteinase 3), *FN1* (fibronectin 1), and *VIM* (vimentin), were among the top overexpressed genes in these tumors (Fig. 5A). *SERPINI2* (Serpin Family I Member 2, a tumor suppressor gene downregulated in PDAC and other cancers^31, 32^) and *MIR217* (microRNA mir-217, a potential tumor suppressor gene in PDAC^33^ which targets KRAS expression) were among the lowest under-expressed genes in the OCM tumors (Fig. 5A).

**Fig. 5.**
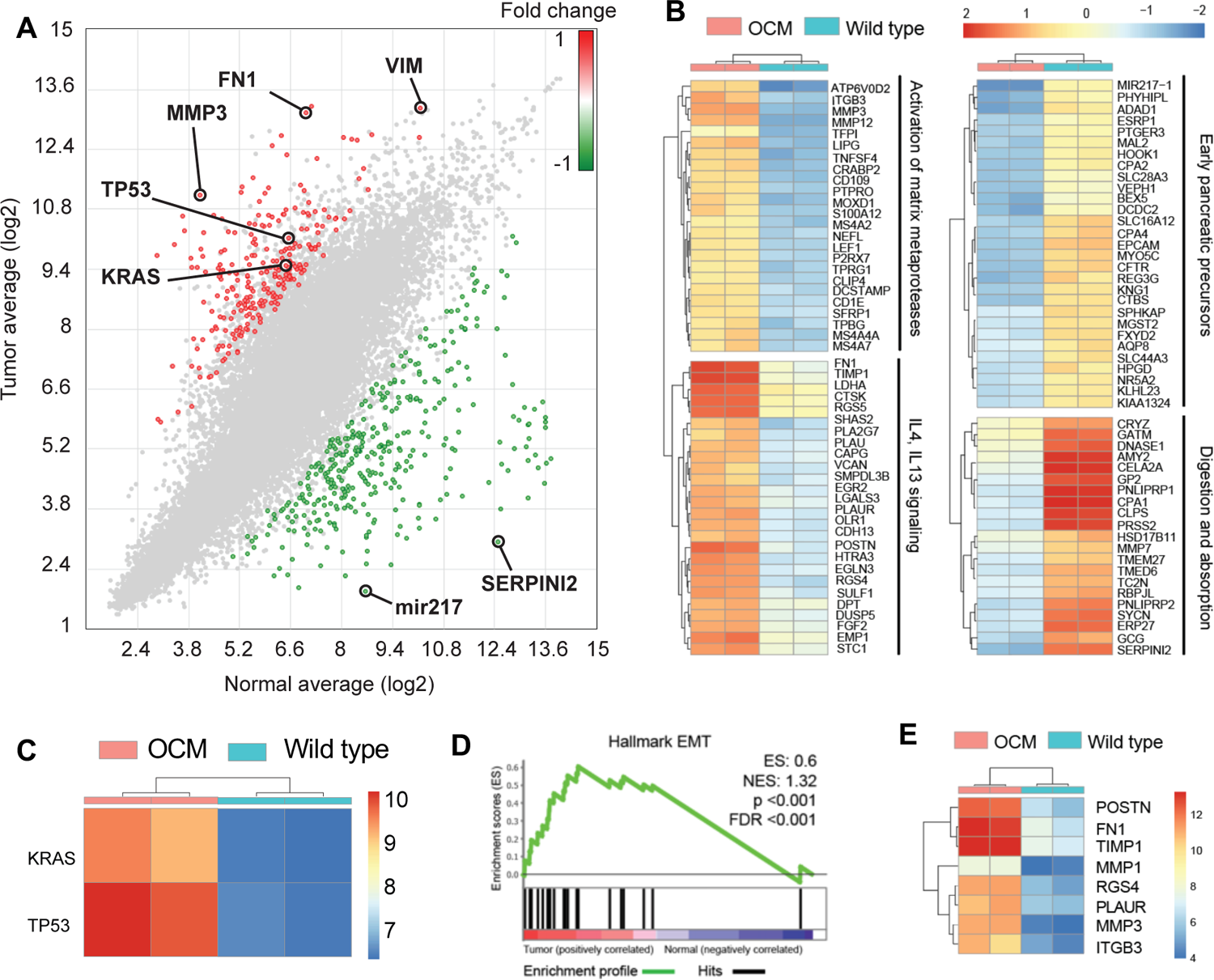
Transcriptomics of OCM pancreatic tumors *vs.* WT pancreas. (A) Differential gene expression scatterplot (RNA-seq). (B) Hierarchical cluster analysis of top 50 upregulated and downregulated genes. (C) *KRAS* and *TP53* expression. (D) Enrichment score for Hallmark EMT genes. (E) Differential expression of select EMT genes.

Hierarchical cluster analysis of the top 50 upregulated and top 50 downregulated genes in OCM pancreatic tumors identified four major biological processes that were altered in the tumors (Fig. 5B): (*i*) matrix metalloproteinases; (*ii*) IL4/IL13 signaling; (*iii*) early pancreatic precursors; and (*iv*) digestion and absorption. Matrix metalloproteases and IL4/IL13 signaling were upregulated, while early pancreatic precursors and digestion/absorption genes were down-regulated (Fig. 5B). Various MMPs and IL4/IL13 signaling are associated with progression of human PDAC.^34–36^ In addition, expression of *KRAS* and *TP53* were upregulated in the OCM pancreatic tumors (Fig. 5C). The Hallmark EMT pathway (widely associated with cancer, including PC^37, 38^) was over-represented in OCM tumors (Fig. 5D). The top enriched (overexpressed) EMT genes in OCM tumors included *MMP1*, *MMP3*, *POSTN* (periostin), *FN1*, *TIMP1* (tissue inhibitor of metalloproteinase 1), *RGS4* (regulator of G-protein signaling), *PLAUR* (plasminogen activator urokinase receptor) and *ITGB3* (integrin subunit beta 3); see Fig. 5E.

## DISCUSSION

This proof-of-principle study demonstrated that tumor could be induced in the pancreas of the *KRAS/TP53* Oncopig rapidly and with reasonable reliability. OCM tumors were predominantly epithelial on histology, less differentiated with respect to gene expression, but still contained mutations and transcriptomes that resembled human PC. However, the fulminant time course (sometimes <2 weeks) with subjects succumbing to a secondary complication of the tumor (pancreatitis and gastric outlet obstruction) may limit the utility of the model in its present form. The intense inflammatory reaction associated with the tumor may have been secondary to an immune response to an acute focal load of neoantigens in these juvenile immunocompetent pigs.

The immune response against tumor induced within OCM skeletal muscle was previously characterized as an intratumoral infiltration of cytolytic CD8β+ T cells,^39^ which presumably was the response observed in this study. Another possible cause of the intense inflammatory reaction may have been a response to adenoviral infection; however, this is unlikely, since none of the WT pigs similarly treated with AdCre developed pancreatitis. One control group not performed in this study was to inject the connecting lobe duct of the Oncopig with the adenoviral vector minus Cre, to determine whether there was atypical OCM response to adenoviral infection not present with WT pigs (admittedly an unlikely possibility).

The clinical course of tumor development and the associated inflammatory reaction might be slowed by decreasing the AdCre dose (with the understanding that lower doses have produced no tumor at all). Another option might be to use a different viral vector (e.g., lentivirus), which might result in lower levels of Cre expression. The anti-tumor lymphocytic inflammatory response also might be mitigated with administration of immunosuppression. However, use of immunosuppression would increase the model complexity and might impact model relevance. The fields of tumor immunomodulation and tumor immunoediting currently are under intense study,^40–43^ and tumor interactions with the immune system in these OCM subjects could be highly relevant.^39^

Co-administration of IL-8 with the AdCre was utilized in this report because of the potential for this chemoattractant to enhance tumor development (IL-8 can increase AdCre entry through the apical membrane of epithelial cells^22, 23^). While this study was not designed and did not have adequate numbers to test IL-8 effect, there may have been a trend for IL-8 enhancement of tumor development; but no firm conclusion drawn. Administration of IL-8 alone did not induce pancreatitis in two control subjects.

Subcutaneous tumor growth has been demonstrated in a proof-of-principle study with implantation of human PC cells (Panc01) into the ears of immunodeficient transgenic pigs (*RAG2*/*IL2RG* deficient).^10^ While immunodeficient orthotopic xeno/allograft models have an advantage with respect to xenografting human PC, there are two issues with this approach: (*i*) the lack of a functional immune system raises issues of clinical relevance (similar to critiques of oncologic studies using immunodeficient mice,^3, 43, 44^); and (*ii*) husbandry of immunodeficient pigs is complex relative to immunocompetent pigs. Hence it has been our preference to develop an immunocompetent porcine model of PC.

Induction of neoplasia with injection of AdCre into the main pancreatic duct of the Oncopig has been described,^11^ but this was a proof-of-principal demonstration in one subject that had microscopic changes only, observed one year after induction. Another group was able to percutaneously access the Oncopig pancreas under CT-guidance, and was able to induce tumor that became grossly evident within a month.^9^ While the percutaneous method avoids a laparotomy, it does not restrict or control the transformation events. This results in nonspecific transformation of many cell types, resulting in pleomorphic tumors.

Tumor pleomorphism has been a described consequence of nondirected injection of AdCre into OCM tissue.^9, 11, 45^ While it is clear that AdCre injection into the Oncopig at various sites will produce neoplasia, the clinical relevance of resultant pleomorphic tumors is questionable. Our OCM pancreatic tumor induction technique is intended to avoid this nonspecific transformation by directing injection of AdCre into the duct of the surgically isolated connecting lobe, which should produce more specific, epithelial transformation. However, we cannot rule out the possibility that OCM pancreatic tumor in this study consisted of multiple transformed cell types.

Exomic analysis of the OCM pancreatic tumors confirmed the presence of the *KRAS*^G12D^ and *TP53*^R167H^ transgenes, along with additional mutations in *KRAS*, *TP53*, and *SMAD4* in some of the OCM subjects (2/4, 3/4 and 1/4 Oncopigs, respectively). In human PDAC, mutations in *KRAS*, *TP53*, and *SMAD4* occur in 74, 53, and 23% of patients, respectively (Table S3). The small number of OCM tumors (N = 4), however, did not permit formal comparison of mutational rates between humans and Oncopigs. Transcriptomic analysis confirmed expression of the *KRAS*^G12D^ and *TP53*^R167H^ transgenes in OCM tumors, along with an increased proliferative profile and expression of acidic mucins. Moreover, the EMT profile and IL4/IL13 signaling were upregulated in these tumors. Human PDAC has a fibroinflammatory tumor microenvironment, with high levels of mucins, interleukins and EMT-associated TIMPs,^34, 35, 37, 38^ which all are associated with PC progression and metastasis.^36, 46^ OCM tumors also had less expression of pancreatic precursor and digestion/absorption genes, suggesting some tumor dedifferentiation.

The presence of regional surface (“drop”) metastases in the OCM subjects suggested that intraperitoneal spillage of the AdCre during the injection procedure may have occurred, with resultant formation of peritoneal tumor nodules. This incidental finding may have implications in the development of an OCM-based model of peritoneal carcinomatosis. With regard to the cytokine arrays, OCM subjects with tumor did show elevation of a cytokine subset, but these relative increases were not significantly different compared to subjects without tumor. This study was limited by availability of OCM litters which resulted in the unbalanced OCM male:female ratio (Table 1).

## Financial Support

this work was funded by grants from the University of Nebraska Medical Center, the Buffett Cancer Center, and the National Cancer Institute.

## Conflict of interest statement

none of the authors have any relevant conflicts of interest to declare.

## Author contributions

Data availability statement: Gene expression microarray data is available at GEO with accession number: GSE203011. Exome sequencing data is available at SRA with accession number: PRJNA838612.

## Supplemental Material

### Supplemental Figures

**Fig. S1** Genotyping

**Fig. S2** Supplemental Methods

**Fig. S3** Preliminary OCM Studies

**Fig. S4** Additional OCM Tumor Images

**Fig. S5** Cytokine Plots

**Fig. S6** ARRIVE Information

**Fig. S7** Author Contributions

### Supplemental Tables

**Table S1** Porcine Descriptive Data

**Table S2** Cytokine Data

**Table S3** Variations in Major PC Genes Within OCM Tumors

**Table S4** Variations in PC-Related Genes Within OCM Tumors

**Table S5** Alteration Frequency of *KRAS*, *TP53*, and *SMAD4* in Human PC

**Table S6** Antibody Information

## Notes

### Competing Interest Statement

The authors have declared no competing interest.

### Summary of Updates

Includes additional IHC and transcriptome data, updated Discussion and author list

