## Supplementary figures and images for "Induction of pancreatic neoplasia in the *KRAS*/*TP53* Oncopig"

### Supplemental Figure 1

#### Slide 1
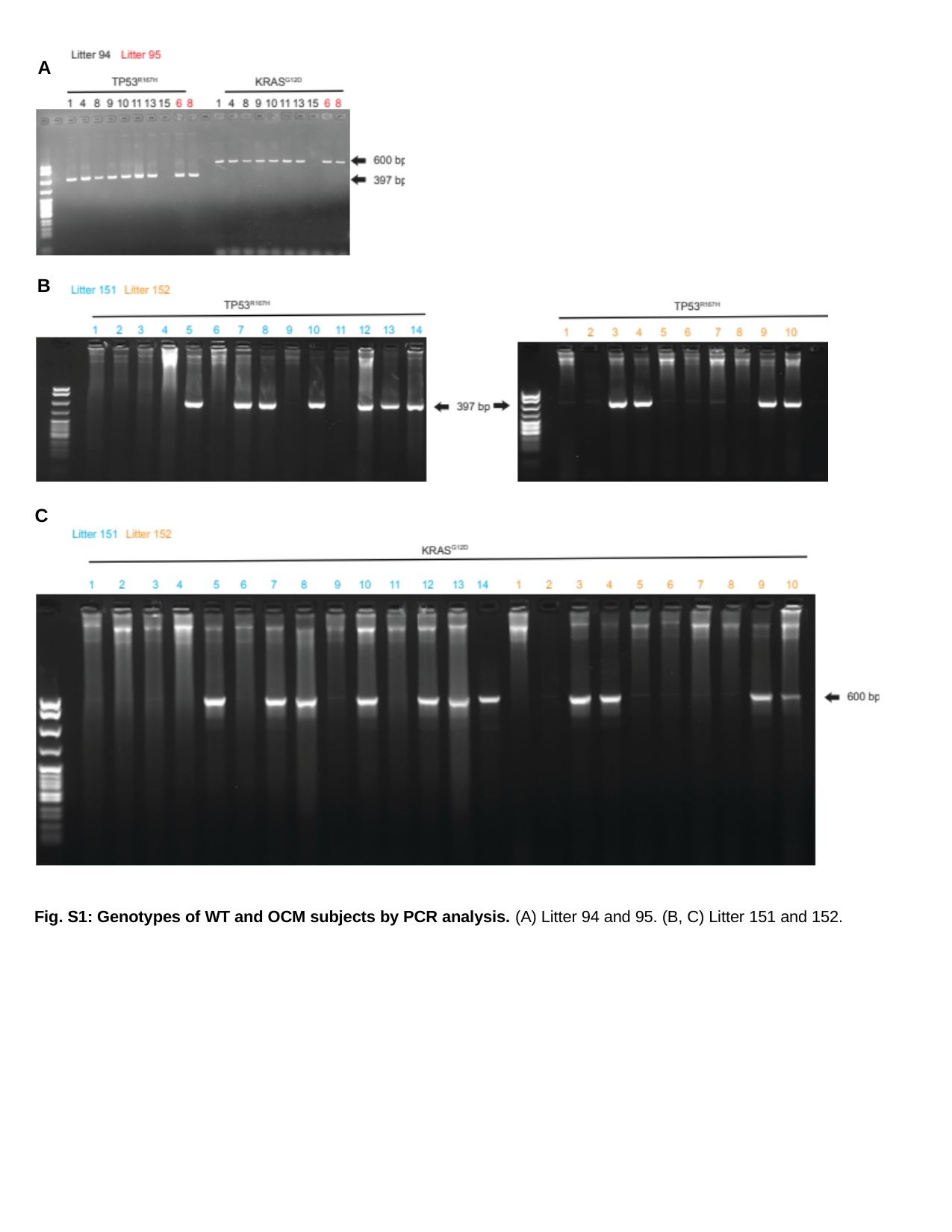

A
B
C
Fig. S1: Genotypes of WT and OCM subjects by PCR analysis. (A) Litter 94 and 95. (B, C) Litter 151 and 152.
