## Supplemental Figure 2 for "Induction of pancreatic neoplasia in the *KRAS*/*TP53* Oncopig"

**Fig. S2.** Supplemental Methods.

***Survival Procedure: Set-Up and Anesthesia***

Swine were fasted for 24 h prior to the procedure, with free access to water. On day zero, subjects were weighed, and underwent induction with ketamine (2.2 mg/kg), Telazol® (1:1 w/w tiletamine:zolazepam, 4.4 mg/kg) and xylazine (2.2 mg/kg), given as a single IM (intra-muscular) injection. Buprenorphine SR (0.2 mg/kg) was given as an IM injection into the right hip. EKG, pulse oximetry, and lingual end-tidal CO_2_ monitors were placed and connected to a BM5 Bionet monitor (bionetus.com). Endotracheal intubation was performed with a 6-7 mm ID (internal diameter) tube. The subject rested on a water-circulated warming blanket that was set at 102˚F. An auricular IV (intravenous) line was placed, and LR (Lactated Ringers solution) was administered at 500 mL/h. Cefovecin sodium (Convenia®; 8 mg/kg IM) and Buprenorphine SR (0.2mg/kg SC or subcutaneous) were administered during the induction period. Anesthesia was maintained with isoflurane (1-2%) and supplemental oxygen (3-5 L/min) using a Matrx® ventilator (midmark.com). The ventilator rate initially was set at 12-15 breaths per minute with a tidal volume of 8 mL/kg, and subsequently adjusted to maintain the EtCO2 at 40-50 mm Hg. Phlebotomy was performed on a forelimb or auricular vein. The ventral abdomen, groins, and thorax were scrubbed with chlorhexidine soap, and then depilated with electric clippers. Cotton blankets were placed over non-surgical areas to minimize subject heat loss. The final abdominal preparation was performed with alcohol-based chlorhexidine (ChloraPrep™; bd.com), and then the upper midline region was sterilely draped.

***Reagents***

Ad5CMVCre-eGFP (AdCre) was purchased from the University of Iowa Vector Core (vector-core.medicine.uiowa.edu). Porcine IL-8 was purchased from Novus Biological (NBP2-35234; novusbio.com). General chemicals were purchased from Millipore Sigma (www.sigmaaldrich.com).

***Standards, Rigor, Reproducibility, and Transparency***

To the extent possible, the animal studies of this report were designed, performed, and reported in accordance with both the ARRIVE recommendations (Animal Research: Reporting of *In Vivo* Experiments; see Fig. S6)^35^ and the National Institutes of Health Principles and Guidelines for Reporting Preclinical Research;^36^ for details and exceptions, refer to Supplemental Information.

***Animal Welfare Statement***

The animals utilized for this report were maintained and treated in accordance with the *Guide for the Care and Use of Laboratory Animals* (8^th^ ed.)^37^ and in accordance with the Animal Welfare Act of the United States (U.S. Code 7, Sections 2131 – 2159). The animal protocol pertaining to this manuscript was approved by the Institutional Animal Care and Use Committee (IACUC) of the VA Nebraska-Western Iowa Health Care System (ID number 1124) and by the IACUC of the University of Nebraska Medical Center (ID number 19-053-FC). All procedures were performed in animal facilities approved by the Association for Assessment and Accreditation of Laboratory Animal Care International (AAALAC; www.aaalac.org) and by the Office of Laboratory Animal Welfare of the Public Health Service (grants.nih.gov/grants/olaw/olaw.htm). All surgical procedures were performed under isoflurane anesthesia, and all efforts were made to minimize suffering. Euthanasia was performed in accordance with the AVMA Guidelines.^38^

***Tissue processing, Histology and Immunohistochemistry***

Tissue was either formalin fixed and sent for further processing, or fresh tissue was taken and digested for *in vitro* analysis (see below). Tissue that was cut from formalin-fixed samples was stained with H&E for pathological examination. IHC was performed by deparaffinizing slides and heating using citric acid antigen retrieval buffer (Vector laboratories, H3300). Following antigen retrieval, endogenous peroxidases were quenched with 3% hydrogen peroxide solution for 5 min. Blocking was done using 2.5% goat serum blocking buffer. Primary antibodies were incubated overnight at 4ºC and are listed in Table S6. Vector Laboratories (vectorlabs.com) ImmPress® goat anti-mouse (MP-7452) or anti-rabbit (MP-7451) IgG polymer kits were used for primary antibody detection per the manufacturer’s instructions. Detection of the HRP/ peroxidase enzyme was performed with SignalStain® DAB Substrate Kit from Cell Signaling (cat. no. 8059; www.cellsignal.com), per the manufacturer’s instructions. Alcian blue staining was performed using the Alcian Blue stain kit (pH 2.5, Mucin Stain) from abcam (cat. no. ab150662; www.abcam.com). For each slide, four different images were captured at 20X or 40X magnification and quantification was performed using open source software for digital image analysis (ImageJ; imagej.nih.gov/ij).

***Cytokine Analysis***

Blood was extracted with a syringe and needle before surgery and after necrops, and then immediately transferred to 10 mL heparinized tubes. Plasma was extracted per the Eve Technologies sample preparation protocol (evetechnologies.com; see Supplemental Information). Samples were sent to Eve Technologies for performance of a 13-plex (PD13) cytokine/chemokine array, which evaluated 13 different markers (IL-1α, IL-1β, IL-1ra, IL-2, IL-4, IL-6, IL-8, IL-10, IL-12, IL-18, GM-CSF, IFNγ, TNFα) using the Millipore MILLIPLEX MAP Porcine Cytokine/Chemokine Magnetic Bead Panel (cat. no. PCYTMG-23K-13PX, www.merckmillipore.com). Samples were detected through a multiplexed immunoassay that was analyzed with a BioPlex 200 (Bio-Rad, www.bio-rad.com).

***Exome sequencing and analysis***

Exome sequences were captured from tumor genomic DNA (isolated by AllPrep DNA/RNA Mini Kit, Cat. No. 80204, www.qiagen.com) by SeqCap EZ HyperCap Exome capture probes, designed for pigs (Roslin Pig Exome Design Version 1) according to manufacture protocol (Roche Sequencing and Life Science, Wilmington, MA, USA), and sequenced on a NextSeq500 instrument (Illumina Inc., San Diego, CA, USA). The sequences were merged and trimmed using the fqtirm tool (https://ccb.jhu.edu/software/fqtrim) to remove adapters, terminal unknown bases (Ns), and low quality 3’ regions (Phred score < 30). Sequences then were processed by bcbio-nextgen 1.2.4 (https://doi.org/10.5281/zenodo.3564938) tool kit with QC (quality control) check by MultiQC 1.9, alignment by bwa, and variant calling by FreeBayes, GATK-HaplotypeCaller, and SAMtools. Variants were called ≥2 among the three variant callers to generate the final VCF (variant call format) file. The bcftool in SAMtools was used to generate a single common VCF file for each comparison group and was also used to generate a unique VCF file for each sample in each comparison. The VCF files were then subjected to VEP (variant effect predictor) tool to annotate and predict the effects of each variant. Finally, Circos was used to visualize the variants on pig reference genome. The exome sequencing data is available at SRA with accession number: PRJNA838612.

***Transcriptomic analysis***

Total RNA was isolated from normal pig pancreas and pancreatic tumor samples using AllPrep DNA/RNA Kits (Qiagen, Germantown, MD, USA), analyzed for RNA integrity score (RIN) by TapeStation System (Agilent, Santa Clara, CA, USA), and then hybridized on a Porcine Gene 1.1 ST array (Thermo Fisher Scientific, Waltham, MA, USA). The data was quality checked, normalized, and analyzed on a Transcriptome Analysis Console (Agilent, Santa Clara, CA, USA). GESA (Gene Set Enrichment Analysis) was used to identify enriched expression signatures in the tumor samples. The gene expression microarray data is available at GEO with accession number: GSE203011.

***Eve Technologies serum sample preparation protocol***

1. Allow blood to clot for 30min or more at room temp. After clotting, centrifuge at 1000 x g for 10min at 4°C.
2. Aliquot serum immediately into a pyrogen/endotoxin-free polypropylene tube.
3. Store samples at ≤-20°C (for ~1 month storage life) or ≤-70°C (for more than 1 month storage).
4. Avoid using hemolyzed or lipemic sera.
5. Avoid multiple freeze/thaw cycles (>2 cycles).

***Study Termination and Euthanasia***

The prescribed post-induction observation period was three months. Criteria for early removal from the study and euthanasia were symptoms of failure to thrive (anorexia, lethargy, decreased movement, abnormal breathing, or other signs of distress) or sepsis (fever, wound disruption or drainage). At the time of euthanasia, each subject received sedation with an IM shot of ketamine/Telazol/xylazine, as described above, and then were endotracheally intubated. Inhalational isoflurane (5%) was administered via the ventilator. The prior midline incision was reopened. Inferiorly this incision was extended in paramedian fashion to avoid midline structures (such as the urethra in males). The completed necropsy incision extended from xiphoid process to the pelvic inlet. The bilateral thoracic cavity was entered by transversely incising the diaphragm just inferior to the xiphoid process. The intrathoracic portion of the inferior vena cava was easily identified as it emerged from the liver in the posterior mediastinum. Phlebotomy was performed from the cava, and then Fatal-Plus® (pentobarbital sodium, 390 mg/mL; 1 mL per 4.5 kg body weight) was administered by caval injection. Two minutes after administration of Fatal-Plus®, the inferior vena cava was transected just above the diaphragm to exsanguinate the subject. A gross necropsy involving the lungs, heart, liver, kidneys, pancreas, intestines, bladder, and associated peritoneal surfaces then was performed, with biopsy of any suspicious lesions.

***Statistical Analysis***

Data are reported as mean ± standard deviation. Comparison of means was performed using t-testing or two-way ANOVA as indicated, with the level of significance set at p <0.05.
