## Supplemental Figure 3 for "Induction of pancreatic neoplasia in the *KRAS*/*TP53* Oncopig"

### Slide 1
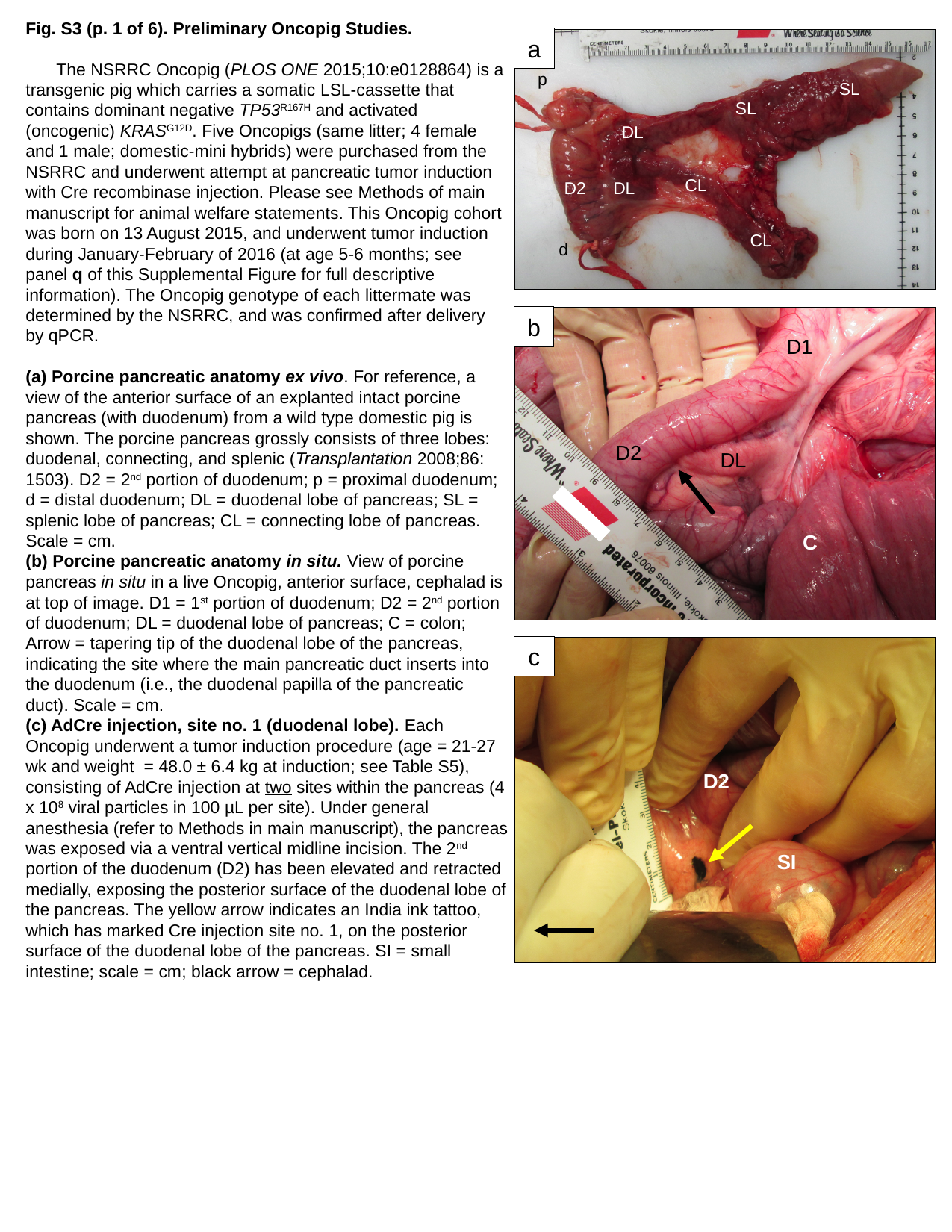

Fig. S3 (p. 1 of 6). Preliminary Oncopig Studies.
	The NSRRC Oncopig (PLOS ONE 2015;10:e0128864) is a transgenic pig which carries a somatic LSL-cassette that contains dominant negative TP53^R167H^ and activated (oncogenic) KRAS^G12D^. Five Oncopigs (same litter; 4 female and 1 male; domestic-mini hybrids) were purchased from the NSRRC and underwent attempt at pancreatic tumor induction with Cre recombinase injection. Please see Methods of main manuscript for animal welfare statements. This Oncopig cohort was born on 13 August 2015, and underwent tumor induction during January-February of 2016 (at age 5-6 months; see panel q of this Supplemental Figure for full descriptive information). The Oncopig genotype of each littermate was determined by the NSRRC, and was confirmed after delivery by qPCR.
(a) Porcine pancreatic anatomy ex vivo. For reference, a view of the anterior surface of an explanted intact porcine pancreas (with duodenum) from a wild type domestic pig is shown. The porcine pancreas grossly consists of three lobes: duodenal, connecting, and splenic (Transplantation 2008;86: 1503). D2 = 2^nd^ portion of duodenum; p = proximal duodenum; d = distal duodenum; DL = duodenal lobe of pancreas; SL = splenic lobe of pancreas; CL = connecting lobe of pancreas. Scale = cm.
(b) Porcine pancreatic anatomy in situ. View of porcine pancreas in situ in a live Oncopig, anterior surface, cephalad is at top of image. D1 = 1^st^ portion of duodenum; D2 = 2^nd^ portion of duodenum; DL = duodenal lobe of pancreas; C = colon; Arrow = tapering tip of the duodenal lobe of the pancreas, indicating the site where the main pancreatic duct inserts into the duodenum (i.e., the duodenal papilla of the pancreatic duct). Scale = cm.
(c) AdCre injection, site no. 1 (duodenal lobe). Each Oncopig underwent a tumor induction procedure (age = 21-27 wk and weight = 48.0 ± 6.4 kg at induction; see Table S5), consisting of AdCre injection at two sites within the pancreas (4 x 10^8^ viral particles in 100 µL per site). Under general anesthesia (refer to Methods in main manuscript), the pancreas was exposed via a ventral vertical midline incision. The 2^nd^ portion of the duodenum (D2) has been elevated and retracted medially, exposing the posterior surface of the duodenal lobe of the pancreas. The yellow arrow indicates an India ink tattoo, which has marked Cre injection site no. 1, on the posterior surface of the duodenal lobe of the pancreas. SI = small intestine; scale = cm; black arrow = cephalad.
a
p
SL
SL
DL
CL
D2
DL
CL
d
b
D1
D2
DL
C
c
D2
SI

### Slide 2
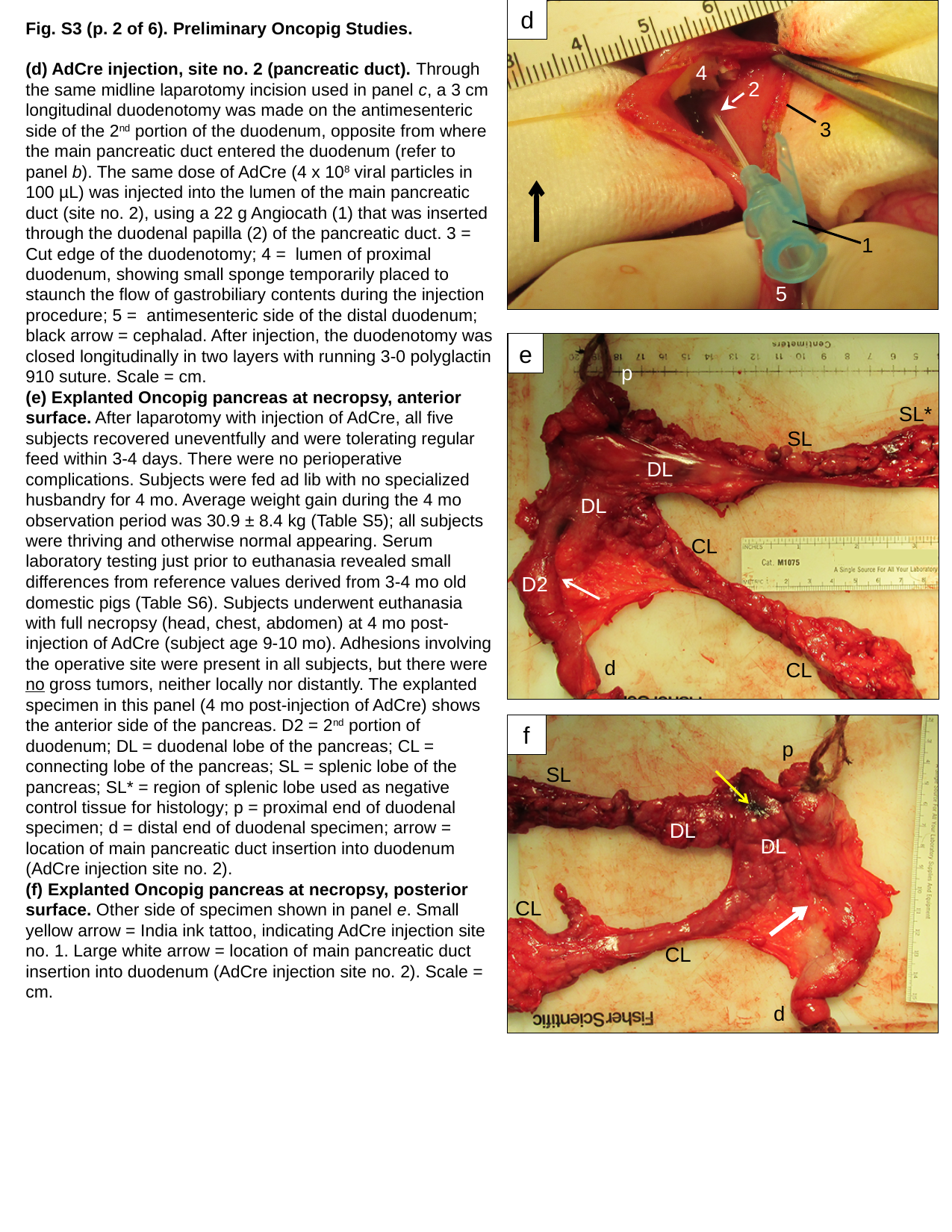

d
Fig. S3 (p. 2 of 6). Preliminary Oncopig Studies.
(d) AdCre injection, site no. 2 (pancreatic duct). Through the same midline laparotomy incision used in panel c, a 3 cm longitudinal duodenotomy was made on the antimesenteric side of the 2^nd^ portion of the duodenum, opposite from where the main pancreatic duct entered the duodenum (refer to panel b). The same dose of AdCre (4 x 10^8^ viral particles in 100 µL) was injected into the lumen of the main pancreatic duct (site no. 2), using a 22 g Angiocath (1) that was inserted through the duodenal papilla (2) of the pancreatic duct. 3 = Cut edge of the duodenotomy; 4 = lumen of proximal duodenum, showing small sponge temporarily placed to staunch the flow of gastrobiliary contents during the injection procedure; 5 = antimesenteric side of the distal duodenum; black arrow = cephalad. After injection, the duodenotomy was closed longitudinally in two layers with running 3-0 polyglactin 910 suture. Scale = cm.
(e) Explanted Oncopig pancreas at necropsy, anterior surface. After laparotomy with injection of AdCre, all five subjects recovered uneventfully and were tolerating regular feed within 3-4 days. There were no perioperative complications. Subjects were fed ad lib with no specialized husbandry for 4 mo. Average weight gain during the 4 mo observation period was 30.9 ± 8.4 kg (Table S5); all subjects were thriving and otherwise normal appearing. Serum laboratory testing just prior to euthanasia revealed small differences from reference values derived from 3-4 mo old domestic pigs (Table S6). Subjects underwent euthanasia with full necropsy (head, chest, abdomen) at 4 mo post-injection of AdCre (subject age 9-10 mo). Adhesions involving the operative site were present in all subjects, but there were no gross tumors, neither locally nor distantly. The explanted specimen in this panel (4 mo post-injection of AdCre) shows the anterior side of the pancreas. D2 = 2^nd^ portion of duodenum; DL = duodenal lobe of the pancreas; CL = connecting lobe of the pancreas; SL = splenic lobe of the pancreas; SL* = region of splenic lobe used as negative control tissue for histology; p = proximal end of duodenal specimen; d = distal end of duodenal specimen; arrow = location of main pancreatic duct insertion into duodenum (AdCre injection site no. 2).
(f) Explanted Oncopig pancreas at necropsy, posterior surface. Other side of specimen shown in panel e. Small yellow arrow = India ink tattoo, indicating AdCre injection site no. 1. Large white arrow = location of main pancreatic duct insertion into duodenum (AdCre injection site no. 2). Scale = cm.
4
2
3
1
5
e
p
SL*
SL
DL
DL
CL
D2
d
CL
f
p
SL
DL
DL
CL
CL
d

### Slide 3
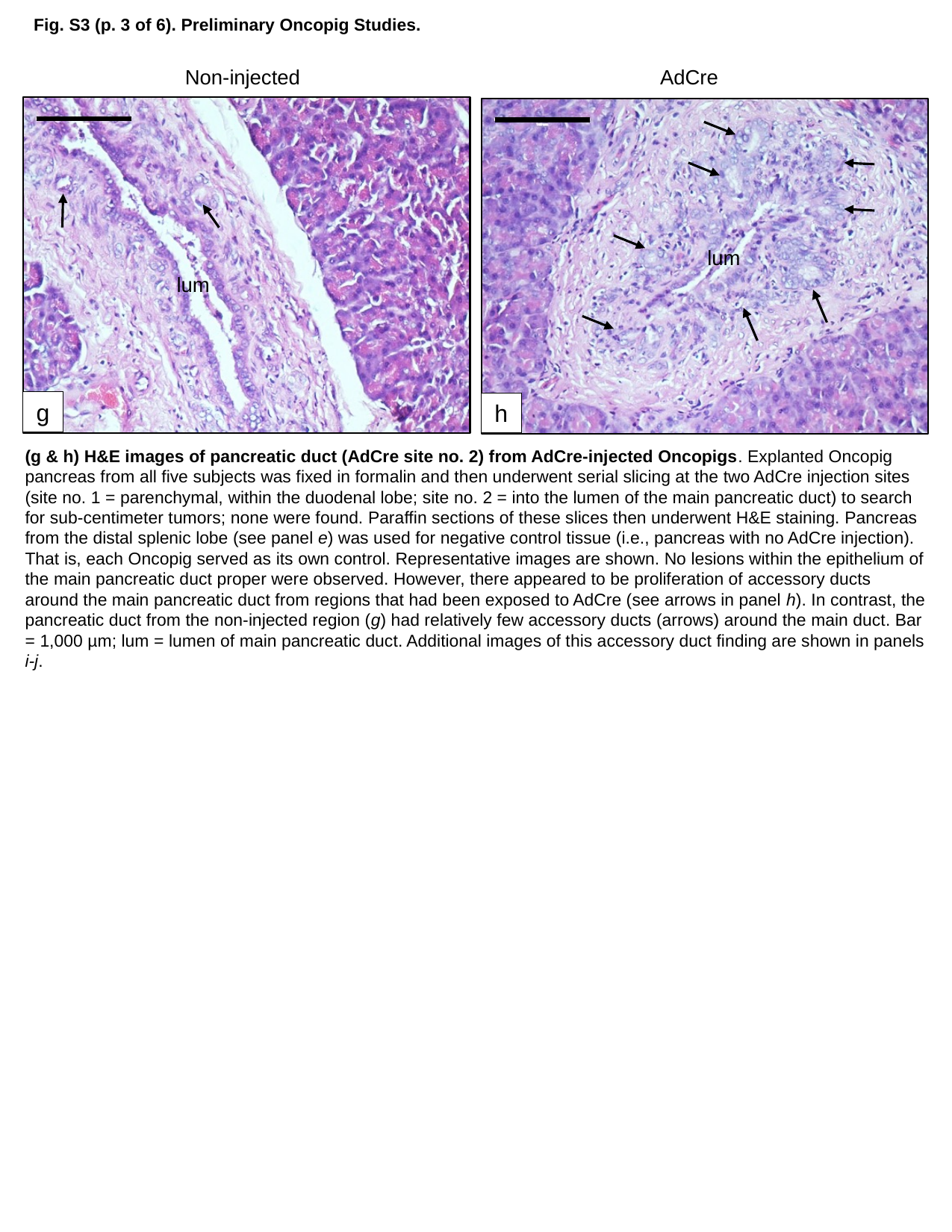

Fig. S3 (p. 3 of 6). Preliminary Oncopig Studies.
Non-injected
AdCre
lum
lum
g
h
(g & h) H&E images of pancreatic duct (AdCre site no. 2) from AdCre-injected Oncopigs. Explanted Oncopig pancreas from all five subjects was fixed in formalin and then underwent serial slicing at the two AdCre injection sites (site no. 1 = parenchymal, within the duodenal lobe; site no. 2 = into the lumen of the main pancreatic duct) to search for sub-centimeter tumors; none were found. Paraffin sections of these slices then underwent H&E staining. Pancreas from the distal splenic lobe (see panel e) was used for negative control tissue (i.e., pancreas with no AdCre injection). That is, each Oncopig served as its own control. Representative images are shown. No lesions within the epithelium of the main pancreatic duct proper were observed. However, there appeared to be proliferation of accessory ducts around the main pancreatic duct from regions that had been exposed to AdCre (see arrows in panel h). In contrast, the pancreatic duct from the non-injected region (g) had relatively few accessory ducts (arrows) around the main duct. Bar = 1,000 µm; lum = lumen of main pancreatic duct. Additional images of this accessory duct finding are shown in panels i-j.

### Slide 4
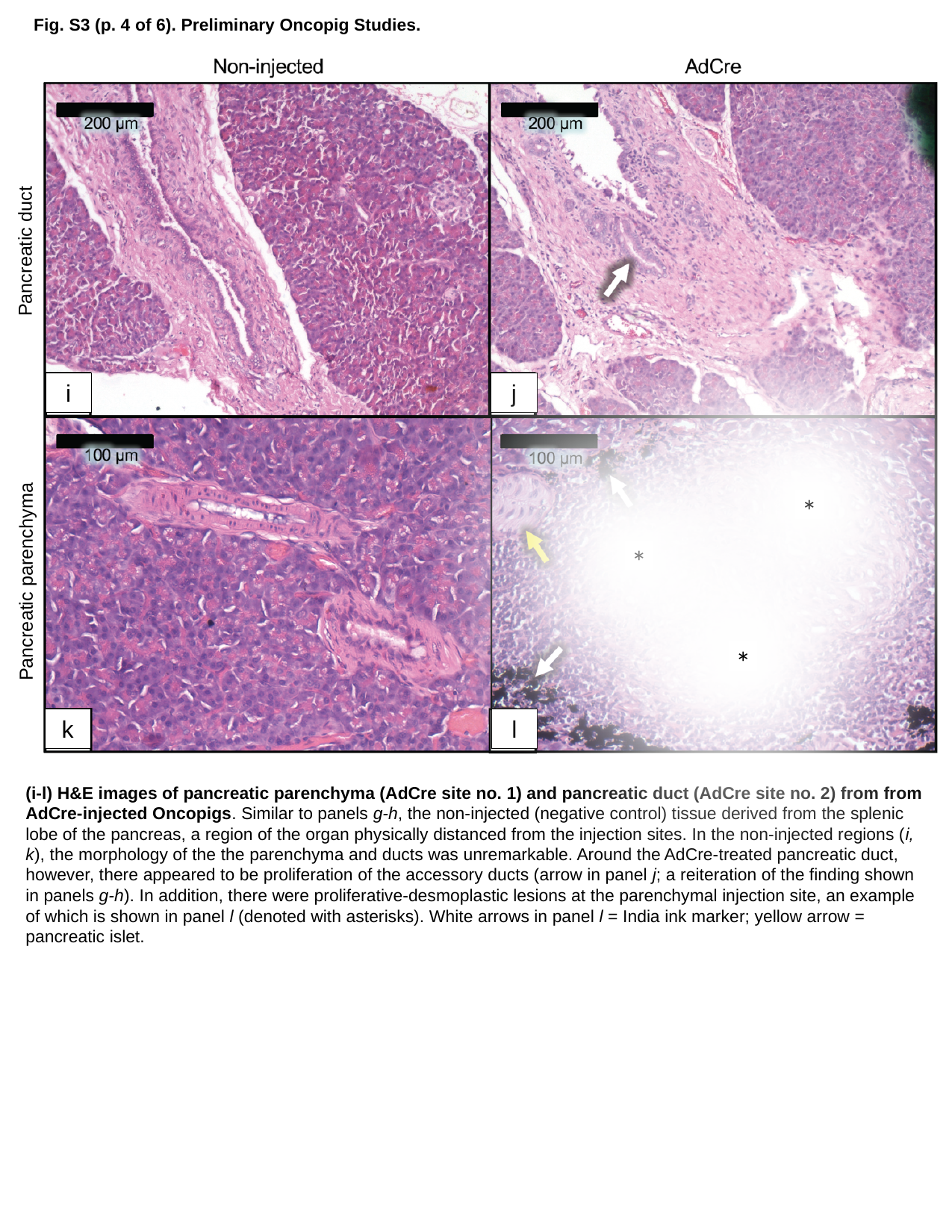

Fig. S3 (p. 4 of 6). Preliminary Oncopig Studies.
Pancreatic duct
j
i
*
*
Pancreatic parenchyma
*
k
l
(i-l) H&E images of pancreatic parenchyma (AdCre site no. 1) and pancreatic duct (AdCre site no. 2) from from AdCre-injected Oncopigs. Similar to panels g-h, the non-injected (negative control) tissue derived from the splenic lobe of the pancreas, a region of the organ physically distanced from the injection sites. In the non-injected regions (i, k), the morphology of the the parenchyma and ducts was unremarkable. Around the AdCre-treated pancreatic duct, however, there appeared to be proliferation of the accessory ducts (arrow in panel j; a reiteration of the finding shown in panels g-h). In addition, there were proliferative-desmoplastic lesions at the parenchymal injection site, an example of which is shown in panel l (denoted with asterisks). White arrows in panel l = India ink marker; yellow arrow = pancreatic islet.

### Slide 5
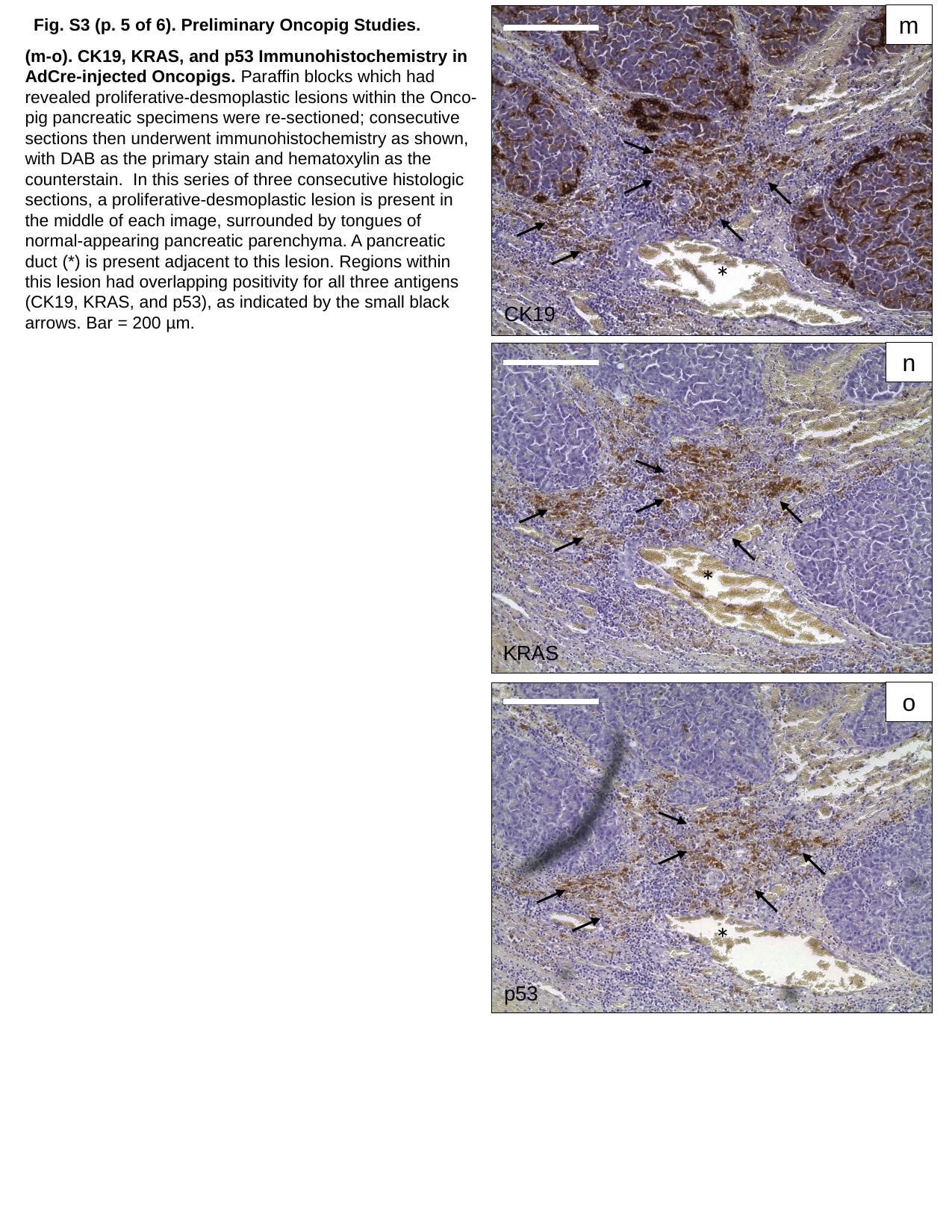

m
Fig. S3 (p. 5 of 6). Preliminary Oncopig Studies.
(m-o). CK19, KRAS, and p53 Immunohistochemistry in AdCre-injected Oncopigs. Paraffin blocks which had revealed proliferative-desmoplastic lesions within the Onco-pig pancreatic specimens were re-sectioned; consecutive sections then underwent immunohistochemistry as shown, with DAB as the primary stain and hematoxylin as the counterstain. In this series of three consecutive histologic sections, a proliferative-desmoplastic lesion is present in the middle of each image, surrounded by tongues of normal-appearing pancreatic parenchyma. A pancreatic duct (*) is present adjacent to this lesion. Regions within this lesion had overlapping positivity for all three antigens (CK19, KRAS, and p53), as indicated by the small black arrows. Bar = 200 µm.
*
CK19
n
*
KRAS
o
*
p53

### Slide 6
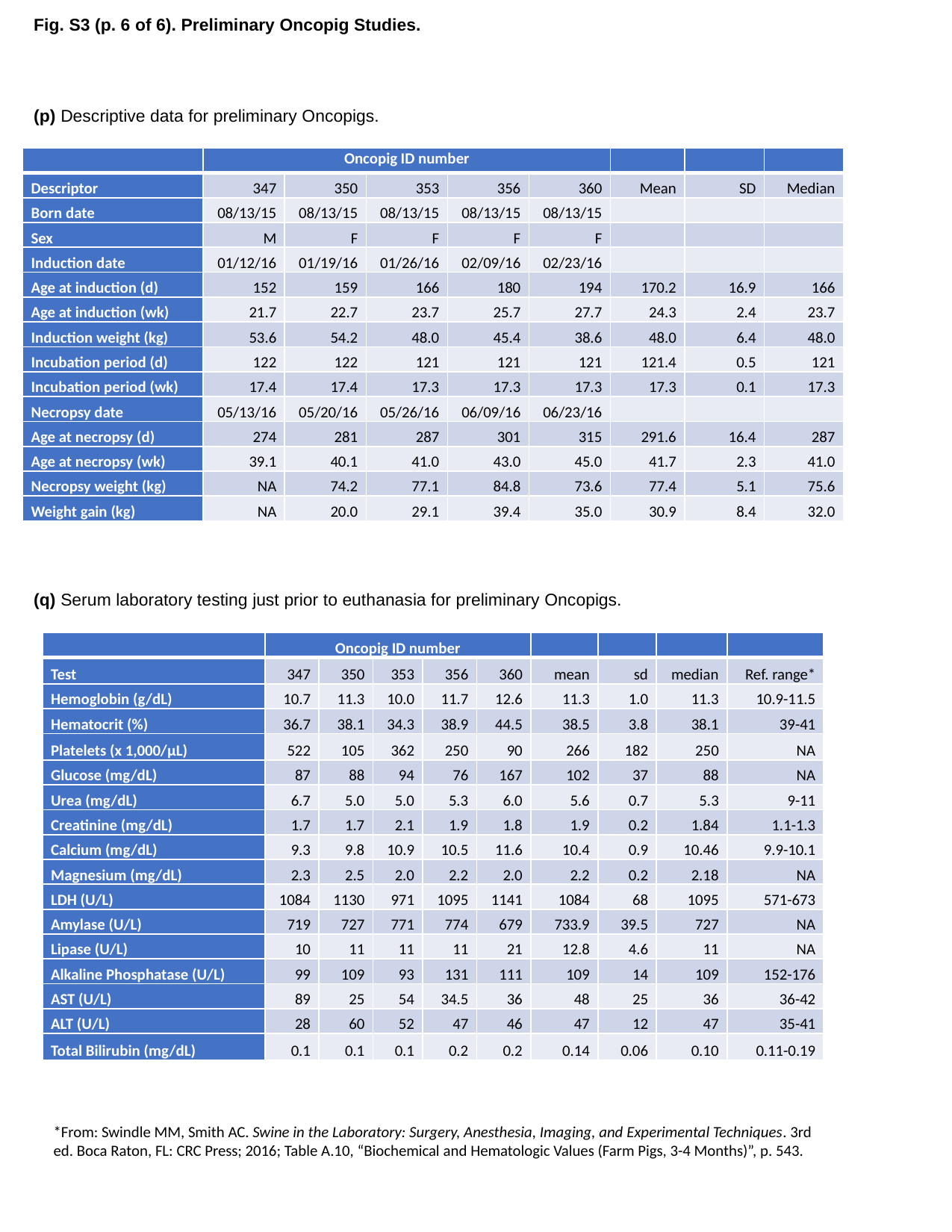

Fig. S3 (p. 6 of 6). Preliminary Oncopig Studies.
(p) Descriptive data for preliminary Oncopigs.
| | Oncopig ID number | | | | | | | |
| --- | --- | --- | --- | --- | --- | --- | --- | --- |
| Descriptor | 347 | 350 | 353 | 356 | 360 | Mean | SD | Median |
| Born date | 08/13/15 | 08/13/15 | 08/13/15 | 08/13/15 | 08/13/15 | | | |
| Sex | M | F | F | F | F | | | |
| Induction date | 01/12/16 | 01/19/16 | 01/26/16 | 02/09/16 | 02/23/16 | | | |
| Age at induction (d) | 152 | 159 | 166 | 180 | 194 | 170.2 | 16.9 | 166 |
| Age at induction (wk) | 21.7 | 22.7 | 23.7 | 25.7 | 27.7 | 24.3 | 2.4 | 23.7 |
| Induction weight (kg) | 53.6 | 54.2 | 48.0 | 45.4 | 38.6 | 48.0 | 6.4 | 48.0 |
| Incubation period (d) | 122 | 122 | 121 | 121 | 121 | 121.4 | 0.5 | 121 |
| Incubation period (wk) | 17.4 | 17.4 | 17.3 | 17.3 | 17.3 | 17.3 | 0.1 | 17.3 |
| Necropsy date | 05/13/16 | 05/20/16 | 05/26/16 | 06/09/16 | 06/23/16 | | | |
| Age at necropsy (d) | 274 | 281 | 287 | 301 | 315 | 291.6 | 16.4 | 287 |
| Age at necropsy (wk) | 39.1 | 40.1 | 41.0 | 43.0 | 45.0 | 41.7 | 2.3 | 41.0 |
| Necropsy weight (kg) | NA | 74.2 | 77.1 | 84.8 | 73.6 | 77.4 | 5.1 | 75.6 |
| Weight gain (kg) | NA | 20.0 | 29.1 | 39.4 | 35.0 | 30.9 | 8.4 | 32.0 |
(q) Serum laboratory testing just prior to euthanasia for preliminary Oncopigs.
| | Oncopig ID number | | | | | | | | |
| --- | --- | --- | --- | --- | --- | --- | --- | --- | --- |
| Test | 347 | 350 | 353 | 356 | 360 | mean | sd | median | Ref. range\* |
| Hemoglobin (g/dL) | 10.7 | 11.3 | 10.0 | 11.7 | 12.6 | 11.3 | 1.0 | 11.3 | 10.9-11.5 |
| Hematocrit (%) | 36.7 | 38.1 | 34.3 | 38.9 | 44.5 | 38.5 | 3.8 | 38.1 | 39-41 |
| Platelets (x 1,000/µL) | 522 | 105 | 362 | 250 | 90 | 266 | 182 | 250 | NA |
| Glucose (mg/dL) | 87 | 88 | 94 | 76 | 167 | 102 | 37 | 88 | NA |
| Urea (mg/dL) | 6.7 | 5.0 | 5.0 | 5.3 | 6.0 | 5.6 | 0.7 | 5.3 | 9-11 |
| Creatinine (mg/dL) | 1.7 | 1.7 | 2.1 | 1.9 | 1.8 | 1.9 | 0.2 | 1.84 | 1.1-1.3 |
| Calcium (mg/dL) | 9.3 | 9.8 | 10.9 | 10.5 | 11.6 | 10.4 | 0.9 | 10.46 | 9.9-10.1 |
| Magnesium (mg/dL) | 2.3 | 2.5 | 2.0 | 2.2 | 2.0 | 2.2 | 0.2 | 2.18 | NA |
| LDH (U/L) | 1084 | 1130 | 971 | 1095 | 1141 | 1084 | 68 | 1095 | 571-673 |
| Amylase (U/L) | 719 | 727 | 771 | 774 | 679 | 733.9 | 39.5 | 727 | NA |
| Lipase (U/L) | 10 | 11 | 11 | 11 | 21 | 12.8 | 4.6 | 11 | NA |
| Alkaline Phosphatase (U/L) | 99 | 109 | 93 | 131 | 111 | 109 | 14 | 109 | 152-176 |
| AST (U/L) | 89 | 25 | 54 | 34.5 | 36 | 48 | 25 | 36 | 36-42 |
| ALT (U/L) | 28 | 60 | 52 | 47 | 46 | 47 | 12 | 47 | 35-41 |
| Total Bilirubin (mg/dL) | 0.1 | 0.1 | 0.1 | 0.2 | 0.2 | 0.14 | 0.06 | 0.10 | 0.11-0.19 |
*From: Swindle MM, Smith AC. Swine in the Laboratory: Surgery, Anesthesia, Imaging, and Experimental Techniques. 3rd ed. Boca Raton, FL: CRC Press; 2016; Table A.10, “Biochemical and Hematologic Values (Farm Pigs, 3-4 Months)”, p. 543.
