## Supplemental Figure 4 for "Induction of pancreatic neoplasia in the *KRAS*/*TP53* Oncopig"

### Slide 1
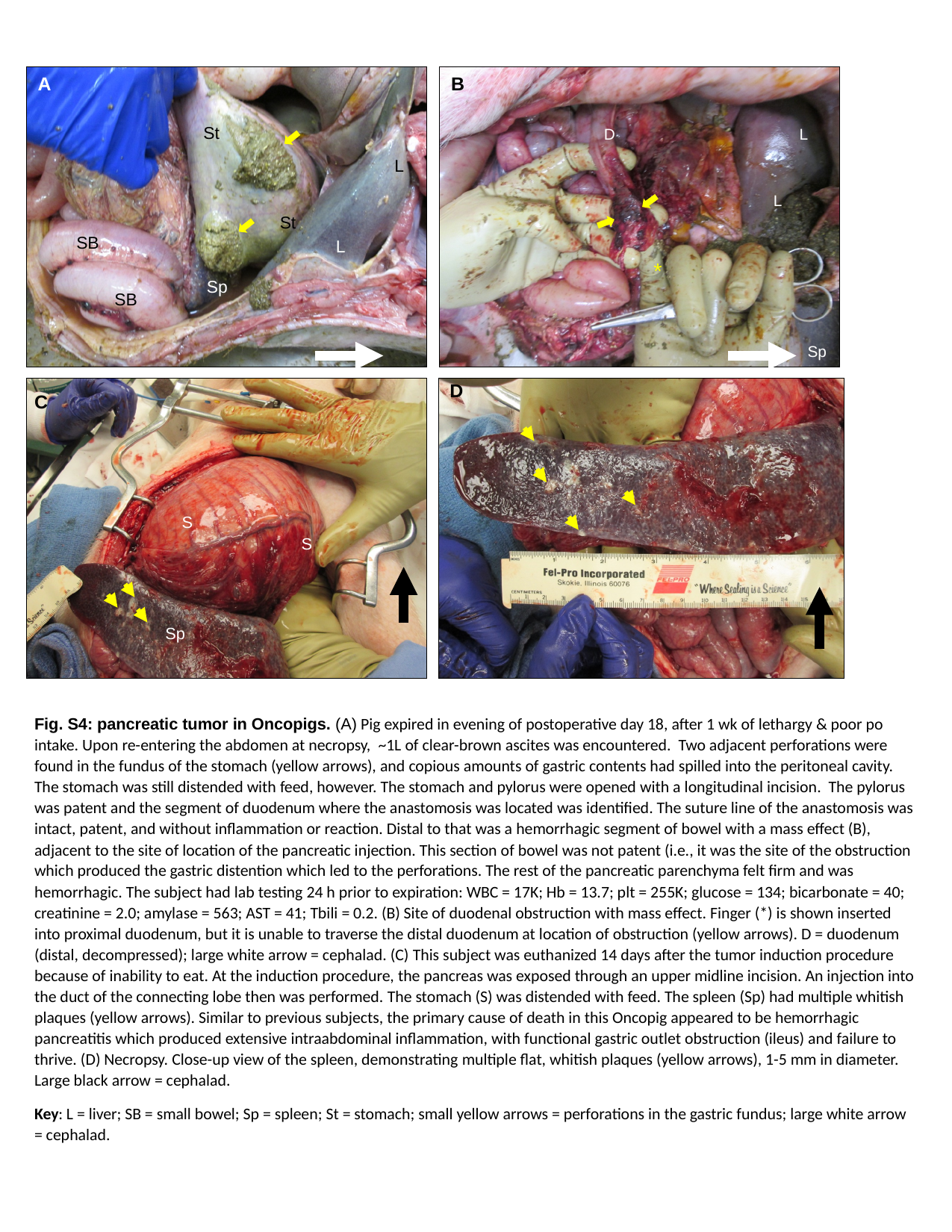

B
A
St
L
St
SB
L
Sp
SB
D
L
L
*
Sp
D
S
S
Sp
C
Fig. S4: pancreatic tumor in Oncopigs. (A) Pig expired in evening of postoperative day 18, after 1 wk of lethargy & poor po intake. Upon re-entering the abdomen at necropsy, ~1L of clear-brown ascites was encountered. Two adjacent perforations were found in the fundus of the stomach (yellow arrows), and copious amounts of gastric contents had spilled into the peritoneal cavity. The stomach was still distended with feed, however. The stomach and pylorus were opened with a longitudinal incision. The pylorus was patent and the segment of duodenum where the anastomosis was located was identified. The suture line of the anastomosis was intact, patent, and without inflammation or reaction. Distal to that was a hemorrhagic segment of bowel with a mass effect (B), adjacent to the site of location of the pancreatic injection. This section of bowel was not patent (i.e., it was the site of the obstruction which produced the gastric distention which led to the perforations. The rest of the pancreatic parenchyma felt firm and was hemorrhagic. The subject had lab testing 24 h prior to expiration: WBC = 17K; Hb = 13.7; plt = 255K; glucose = 134; bicarbonate = 40; creatinine = 2.0; amylase = 563; AST = 41; Tbili = 0.2. (B) Site of duodenal obstruction with mass effect. Finger (*) is shown inserted into proximal duodenum, but it is unable to traverse the distal duodenum at location of obstruction (yellow arrows). D = duodenum (distal, decompressed); large white arrow = cephalad. (C) This subject was euthanized 14 days after the tumor induction procedure because of inability to eat. At the induction procedure, the pancreas was exposed through an upper midline incision. An injection into the duct of the connecting lobe then was performed. The stomach (S) was distended with feed. The spleen (Sp) had multiple whitish plaques (yellow arrows). Similar to previous subjects, the primary cause of death in this Oncopig appeared to be hemorrhagic pancreatitis which produced extensive intraabdominal inflammation, with functional gastric outlet obstruction (ileus) and failure to thrive. (D) Necropsy. Close-up view of the spleen, demonstrating multiple flat, whitish plaques (yellow arrows), 1-5 mm in diameter. Large black arrow = cephalad.
Key: L = liver; SB = small bowel; Sp = spleen; St = stomach; small yellow arrows = perforations in the gastric fundus; large white arrow = cephalad.
