## Supplemental Figure 5 for "Induction of pancreatic neoplasia in the *KRAS*/*TP53* Oncopig"

### Slide 1
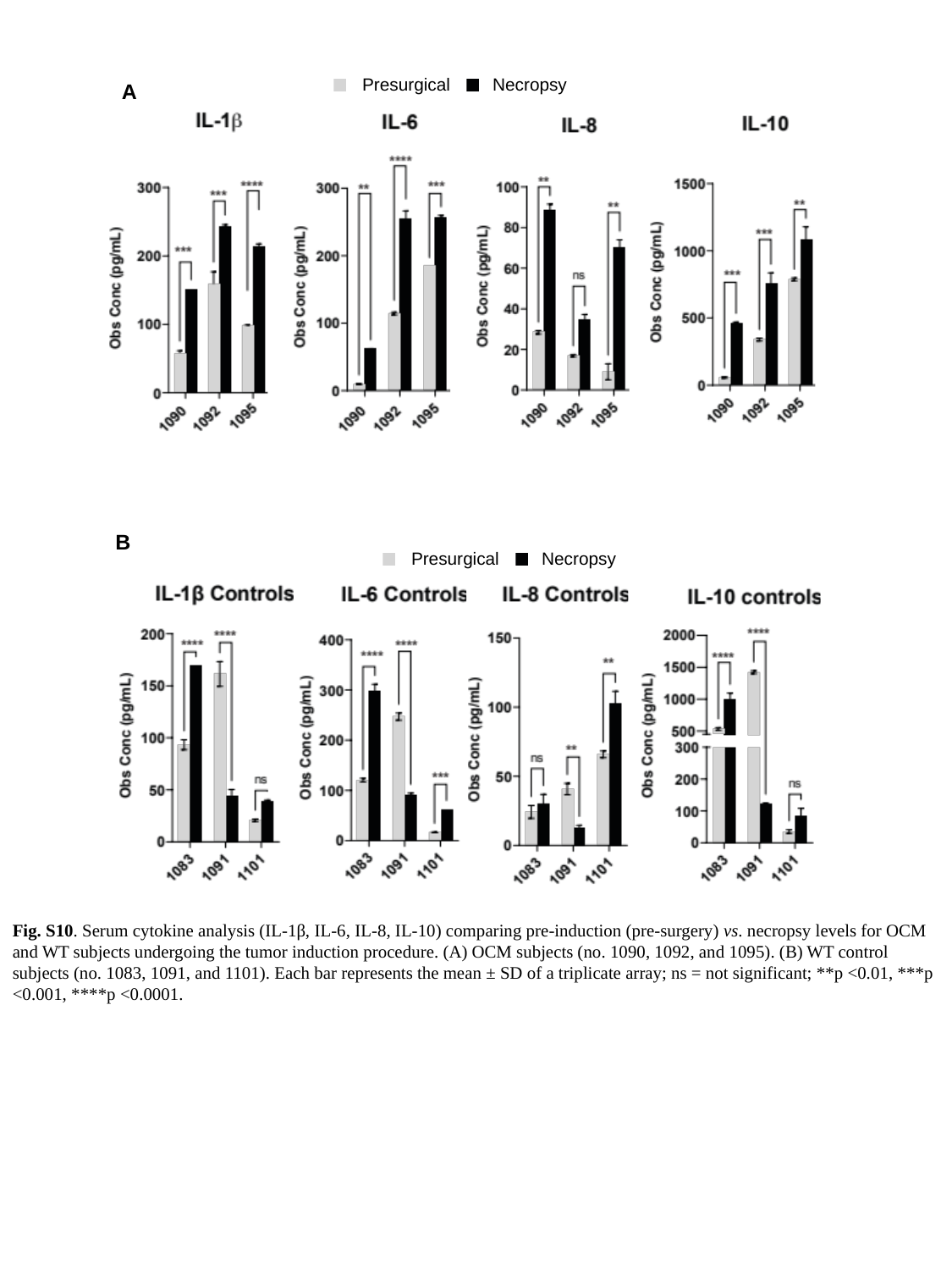

Necropsy
Presurgical
A
B
Necropsy
Presurgical
Fig. S10. Serum cytokine analysis (IL-1β, IL-6, IL-8, IL-10) comparing pre-induction (pre-surgery) vs. necropsy levels for OCM and WT subjects undergoing the tumor induction procedure. (A) OCM subjects (no. 1090, 1092, and 1095). (B) WT control subjects (no. 1083, 1091, and 1101). Each bar represents the mean ± SD of a triplicate array; ns = not significant; **p <0.01, ***p <0.001, ****p <0.0001.
