## Supplemental Figure 7 for "Induction of pancreatic neoplasia in the *KRAS*/*TP53* Oncopig"

**Fig. S7**. Author Contributions [to be completed]

| Contributor Role | Description | Author |
| --- | --- | --- |
| Conceptualization | Ideas; formulation or evolution of overarching research goals and aims. |  |
| Data Curation | Management activities to annotate (produce metadata), scrub data and maintain research data (including software code, where it is necessary for interpreting the data itself) for initial use and later reuse. |  |
| Formal Analysis | Application of statistical, mathematical, computational, or other formal techniques to analyze or synthesize study data. |  |
| Funding Acquisition | Acquisition of the financial support for the project leading to this publication. |  |
| Investigation | Conducting a research and investigation process, specifically performing the experiments, or data/evidence collection. |  |
| Methodology | Development or design of methodology; creation of models. |  |
| Project Administration | Management and coordination responsibility for the research activity planning and execution. |  |
| Resources | Provision of study materials, reagents, materials, patients, laboratory samples, animals, instrumentation, computing resources, or other analysis tools. |  |
| Software | Programming, software development; designing computer programs; implementation of the computer code and supporting algorithms; testing of existing code components. |  |
| Supervision | Oversight and leadership responsibility for the research activity planning and execution, including mentorship external to the core team. |  |
| Validation | Verification, whether as a part of the activity or separate, of the overall replication/reproducibility of results/experiments and other research outputs. |  |
| Visualization | Preparation, creation and/or presentation of the published work, specifically visualization/data presentation. |  |
| Draft Preparation | Creation and/or presentation of the published work, specifically writing the initial draft (including substantive translation). |  |
| Review & Editing | Preparation, creation and/or presentation of the published work by those from the original research group, specifically critical review, commentary or revision – including pre- or post-publication stages. |  |
