## Supplemental Table 3 for "Induction of pancreatic neoplasia in the *KRAS*/*TP53* Oncopig"

**Table S3.** New and existing variations in major pancreatic cancer genes of OCM tumors

| **Genes** | **Variations** | **Location** | **Type** | **Samples** |
| --- | --- | --- | --- | --- |
| KRAS | Deletion (GAT) with rs705858787 | 5:48548834-48548836 | 3 prime UTR variant, cDNA (4665-4667) | 1095, 1096 vs OCM control and Control |
|  | Deletion of rs701486753 (A) | 5:48549683 | Downstream gene variant | 1095, 1096 vs OCM control and Control |
|  | rs329769372 (G/A) | 5:48504818 | Upstream gene variant | 1095 vs OCM control and Control |
|  | Mutation (G/A) | 5:48513419 | Missense variant (G to D) | 1090, 1092, 1095 & 1096 vs Control |
| TP53 | rs336094858 (T/C) | 12:52940975 | Intron variant (T to C) | 1092, 1095 and 1096 vs OCM control |
|  | rs343411452 (T/C) | 12:52942629 | Intron variant (T to C) | 1092, 1095 and 1096 vs OCM control |
|  | rs81211696 (T/G) | 12:52941073 | Synonymous variant (T to G) | 1090, 1092, 1095 & 1096 vs Control |
|  | rs318253531(C/A) | 12:52942206 | Synonymous variant (C to A) | 1090, 1092, 1095 & 1096 vs Control |
|  | rs81211695 (C/T) | 12:52942221 | Synonymous variant (C to T) | 1090, 1092, 1095 & 1096 vs Control |
|  | rs334394956 (G/A) | 12:52942644 | Synonymous variant (G to A) | 1090, 1092, 1095 & 1096 vs Control |
|  | rs324980623 (G/A) | 12:52942707 | Synonymous variant (G to A) | 1090, 1092, 1095 & 1096 vs Control |
|  | rs345021946 (C/A) | 12:52942996 | Synonymous variant (C to A) | 1090, 1092, 1095 & 1096 vs Control |
|  | Mutation (T/C) | 12:52943005 | Intron variant (T to C) | 1090, 1092, 1095 & 1096 vs Control |
|  | Mutation (C/T) | 12:52943212 | Intron variant (C to T) | 1090, 1092, 1095 & 1096 vs Control |
| SMAD4 | Mutation (A/T) | 1:100590467 | Intron variant (A to T) | 1090 vs OCM control and Control |
