## Supplemental Table 4 for "Induction of pancreatic neoplasia in the *KRAS*/*TP53* Oncopig"

**Table S4.** Variations and genetic aberrations in pancreatic cancer related genes in OCM tumors

| **Genes** | **Number of variations** | **Type of variations** | **Samples** |
| --- | --- | --- | --- |
| AKT1 | 15 | SNPs (15), rsID (0) | All samples |
| AKT2 | 12 | SNPs (8), rsID (2) \| single base splice insertion (2) | All samples (SNPs) |
|  |  |  | All vs OCM Control (insertion) |
| AKT3 | 30 | SNPs (24), rsID (24) \| insertions (5), 3 UTR (2), intronic (3) \| six base deletion, intronic, with SNPs (1) | All samples (SNPs) |
|  |  |  | Insertions in 1090, 92 & 96 vs Control and OCM controls |
| ARAF | 3 | SNPs (3), rsID (3) | All vs controls, 1095, 1096 vs OCM controls |
| ARHGEF6 | 63 | SNPs (59), rsID (59) \| one 17bp insertions, intronic (1) \| 3 deletions, downstream (2), intron (1) | All samples (SNPs) |
|  |  |  | Insertion (1096 vs control, OCM control) |
|  |  |  | Deletion (all samples) |
| BAD | 18 | SNPs (18), rsID (17) | All samples |
| BCL2L1 | 3 | SNPs (3), rsID (3) | 1095 vs control, all vs OCM Control |
| BRAF | 34 | SNPs (34), rsID (34) | All samples |
| CASP9 | 106 | SNPs (100), rsID (1) | All samples |
|  |  | 13 bp intronic insertions (1) | Insertions in 1096 vs Control and OCM control |
|  |  | Deletions (4): intronic (3), 3 UTR (1) | Deletions in 1095, 1096 and 1090 |
| CCND1 | 5 | SNP (4), rsID (2) \| Insertion single base intronic (1) | All samples |
|  |  |  | Deletion: 1090, 92, 96 |
| CDK4 | 23 | SNP (23), rsID (21) \| 9 bp insertion intronic (1) | All samples |
|  |  |  | 1095, 1096 vs Control, OCM control |
| CDK6 | 17 | SNP (17), rsID (11) \| single bp deletion (2), double bp deletion (1), 4 bp deletion (1) | All samples |
|  |  |  | 1090, 1095 vs Control and OCM control |
| ERBB2 | 15 | SNPs (13), rsID (13), 5 downstream, 6 intronic, 1 missense, 2 synonymous, 1 upstream \| insertion (2) | All samples |
| RAD51 | 26 | SNPs (26), rsID (25), 11 3 UTR, 10 introns, 1 missense, 1 synonymous, 3 downstream | All samples |
