## Supplemental Table 6 for "Induction of pancreatic neoplasia in the *KRAS*/*TP53* Oncopig"

**Table S6.** Information on antibodies used for immunoblotting and immunohistochemistry.

| **Antigen** | **Product No.** | **Company** | **Website** | **Type** | **Host** | **Clone number** |
| --- | --- | --- | --- | --- | --- | --- |
| Cytokeratin 19 | ab7754 | Abcam | Abcam.com | monoclonal | mouse | A53-B/A2 |
| Mutant KRAS^G12D^ | GTX132407 | Genetex | Genetex.com | polyclonal | rabbit |  |
| Mutant p53 | Bsm-54279R | Bioss | Biossusa.com | monoclonal | rabbit | 2F6 |
| Ki67 | ab16667 | Abcam | Abcam.com | monoclonal | rabbit | SP6 |
| Vimentin | 677801 | Biolegend | Biolegend.com | monoclonal | mouse | O91D3 |
| CD31 | MCA1746GA | Bio-Rad | Bio-rad-antibodies.com | monoclonal | mouse | LCI-4 |
