## Supplemental Table 8 for "Induction of pancreatic neoplasia in the *KRAS*/*TP53* Oncopig"

| **Type of experiment** | | **Number of Oncopigs** | **Number of**  **wild type pigs** |
| --- | --- | --- | --- |
| Control, no Ad-Cre injection | | 2 | 2 |
| Test, Ad-Cre Injection | Main pancreatic duct  (Technique 1 or MPD/T1) | 2 | 0 |
|  | Pancreatic connecting lobe injection  (Technique 2 or CL/T2) | 12 | 4 |
| Transcriptome of the normal porcine pancreas | | 0 | 2 |

**Supplemental Table S8.** Sample size in different experimental groups.
